# Programmable eukaryotic protein expression with RNA sensors

**DOI:** 10.1101/2022.01.26.477951

**Authors:** Kaiyi Jiang, Jeremy Koob, Xi Dawn Chen, Rohan N. Krajeski, Yifan Zhang, Lukas Villiger, Wenyuan Zhou, Omar O. Abudayyeh, Fei Chen, Jonathan S. Gootenberg

**Author notes:** These authors contributed equally (Kaiyi Jiang, Jeremy Koob, Xi Dawn Chen, Rohan N. Krajeski, Yifan Zhang). These authors jointly supervised the work (Omar O. Abudayyeh, Fei Chen, Jonathan S. Gootenberg).

## Abstract

The diversity of cell types and states can be scalably measured and defined by expressed RNA transcripts. However, approaches to programmably sense and respond to the presence of specific RNAs within living biological systems with high sensitivity are lacking. RNA sensors that gate expression of reporter or cargo genes would have diverse applications for basic biology, diagnostics and therapeutics by enabling cell-state specific control of transgene expression. Here, we engineer a novel programmable RNA-sensing technology, Reprogrammable ADAR Sensors (RADARS), which leverages RNA editing by adenosine deaminases acting on RNA (ADAR) to gate translation of a protein payload on the presence of endogenous RNA transcripts. In mammalian cells, we engineer RADARS with diverse payloads, including luciferase and fluorescent proteins, with up to 164-fold activation and quantitative detection in the presence of target RNAs. We show RADARS are functional either expressed from DNA or as synthetic mRNA. Importantly, RADARS can function with endogenous cellular ADAR. We apply RADARS to multiple contexts, including RNA-sensing induced cell death via caspases, cell type identification, and *in vivo* control of synthetic mRNA translation, demonstrating RADARS as a tool with significant potential for gene and cell therapy, synthetic biology, and biomedical research.

**One Sentence Summary:** A new technology utilizing ADAR mediated RNA-editing enables robust reprogrammable protein expression based on target RNA transcripts in mammalian cells, leading to broad applications in basic science research, cell engineering, and gene therapy.

## Main Text

Cell-state specific expression of transgenes is desirable for both fundamental and translational research. For fundamental research, programmable cell-state targeting of transgene expression allows for tracking and perturbing specific cell states to understand their function. Translationally, cell-state sensing and targeting can be critical for cell and gene therapies^1^ and minimally invasive diagnostics^2^. Cell-state targeting is traditionally achieved by either selective transgene delivery or promoter-based control of transgene expression. However, these methods are inflexible and imprecise – and, importantly, do not scale to the diversity of transcriptomically defined cell states. Recent advances in single-cell technologies have revealed a remarkable diversity of cell types and states, specified at the RNA level, that standard viral tropism or cell-specific promoters are not capable of sensing^3,4^. Vectors driven by cell-type specific *cis*-regulatory elements (REs) such as promoters and enhancers improve transgenic specificity^5,6^ but suffer from limitations: a single RE may not define a cell type as REs often act in concert with complex logic^7,8^. Moreover, *de novo* RE development requires additional profiling of active open chromatin and, as they act at the transcriptional level, REs cannot control expression of synthetic mRNA, a valuable asset for mRNA therapeutics and vaccines.

A method that directly and robustly senses and responds to specific RNA markers via simple RNA base pairing rules would provide a direct, programmable path to transcriptional control of transgene expression. Diverse modalities of RNA-based gene control have been developed, including small riboregulators and toehold sensors in bacteria^9^, ribozyme sensors in bacteria^10^, miRNA based synthetic toehold switches^11^, gated Cas9 guide RNAs^12,13^, and mammalian toehold sensors^14^. These approaches, in general, remain difficult to reprogram for new RNAs, have low signal-to-noise in bacteria and mammalian cells, and lack the sensitivity needed to sense endogenous transcripts at different expression levels.

Here, we design RNA-editing based sensors, termed Reprogrammable ADAR Sensors (RADARS), with high reprogrammability, sensitivity, and specificity. RADARS leverages simple base pairing rules and RNA editing to conditionally express an effector protein whenever the target RNA species is detected. The development of RADARS builds upon recent advances in RNA-guided RNA-editing technologies^15–21^ that generate a dsRNA region by hybridizing an exogenous RNA to an endogenous transcript. Engineering an A-C mismatch bubble in this dsRNA region, with a cytosine (C) in the exogenous guide RNA, recruits ADAR to the endogenous transcript for specific editing of the mismatched adenosine (A) to inosine (I), which is translated by the ribosome as a guanine (G) analog^22,23^. RADARS conceptually inverts these RNA-editing systems, directing ADAR to edit an A-C mismatch composed of C on the endogenous target, and A on the exogenously introduced sensor. The targeted adenosine on the sensor lies within a stop codon, which blocks translation of a desired downstream gene payload. Upon sensor binding to the endogenous target transcript, the stop codon is edited and removed, releasing payload expression. Here, we demonstrate that RADARS enable sensitive and specific transgene activation in the presence of RNA target in mammalian cells and tissues, are readily programmable via complementary guides, and can function with endogenous or exogenously provided ADAR, facilitating diverse applications. We perform engineering of RADARS to enable high signal to noise (∼164 fold activation), quantitative detection with high dynamic range, multiplexed transcript detection, diverse effector delivery, detection of endogenous gene perturbations, and *in vivo* activity when delivered as a synthetic mRNA construct.

### Programmable sensing of RNA transcripts with RADARS

We designed RADARS with a simple and modular architecture, consisting of a “guide” region containing one or more hybridization regions complementary to the desired target with an in-frame stop codon and a “payload” region containing the effector protein sequence (Fig. 1a). The UAG stop codon within the guide region is inserted upstream of payload and is designed to oppose a CCA codon, generating the preferred A-C mismatch bubble for adenosine deamination by ADAR. Endogenous ADAR, optionally supplemented with exogenous ADAR, will edit the UAG to UIG in the presence of the complementary target RNA, releasing expression of the payload protein. A constitutive normalizing gene can also be placed upstream of the sensor on the RADAR transcript, or, when RADARS is delivered as a DNA construct, on a separate transcript on the same vector.

**Figure 1:**
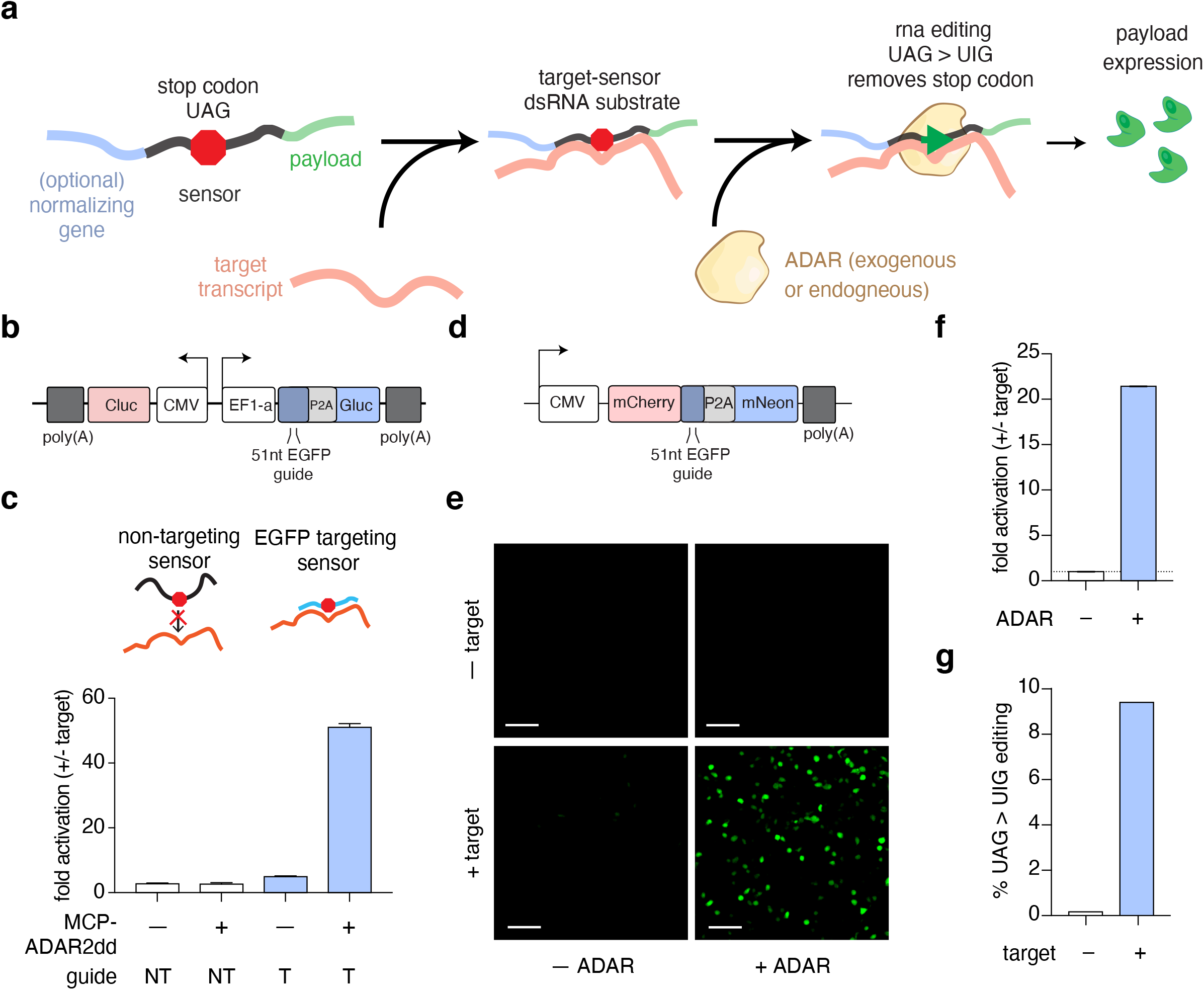
Reprogrammable ADAR Sensors (RADARS) with luciferase and fluorescence outputs can be used to sense reporter transcripts. a) RADARS concept schematic depicting programmable RNA sensing. The sensor RNA contains an optional marker protein, a guide RNA region with a UAG stop (red octagon), and a downstream payload protein. Sensor association with a target RNA forms a duplex with an A-C mismatch, which serves as a substrate for RNA editing by ADAR protein (brown). RNA editing converts the UAG stop codon to UIG, allowing translation of the payload (green protein). b) Schematic for RADARS sensor with a two transcript design containing a payload *Gluc* under RADARS control and a normalizing *Cluc* under constitutive expression. c) Activation of EGFP targeting luciferase RADARS in the presence of target. Ratio fold change is calculated as the ratio of Cluc-normalized sensor payload expression (Gluc/Cluc) to Cluc-normalized payload expression in the absence of target (see Methods). Target is delivered via transfection. The sensor can either recruit endogenous ADAR to sense target EGFP transcript or utilize exogenous delivered ADAR to sense target transcript with enhanced sensitivity. Sensors are evaluated with both EGFP targeting guides (T) and non-targeting scrambled guides (NT) across exogenous MCP-ADAR2dd supplementation and no exogenous ADAR supplementation condition. Data are mean of technical replicates (n= 3) ± s.e.m. d) Schematic showing fluorescent RADARS with a single transcript design, containing a constitutively expressed normalizing fluorescent protein (mCherry) upstream of a RADAR guide controlling mNeon fluorescent protein. e) Images of fluorescence RADARS showing HEK293FT cells expressing payload only in the presence of target, with enhanced activation by the addition of exogenous MCP-ADAR2dd (E488Q, T490A). HEK293FT cells are transfected with combinations of non-expressing EGFP and MCP-ADAR2dd (E488Q, T490A). Top left: sensor; top right: sensor + ADAR; bottom left: sensor + target; bottom right: sensor + target + ADAR. Scale bar, 100 microns. f) Quantification of mNeon fluorescence bulk fold change upon target induction from images in (e); ratio fold change indicates that mNeon/mCherry fluorescence values (fluorescence ratio values) in the presence of target were divided by fluorescence ratio values in the absence of target for ADAR variant (see Methods). Data are mean of technical replicates (n= 3) ± s.d. g) Corresponding editing rate of the sensor UAG codon to UIG in the presence and absence of target. Data obtained via next-generation sequencing of sensor RNA extracted from HEK293FT cells for conditions in (f). Data are mean of technical replicates (n= 3) ± s.d.

To pilot the RADARS concept, we developed a sensor with a 51 nt EGFP transcript-sensing guide and a *Gaussia luciferase* (Gluc) payload. We included a constitutive *Cypridina luciferase* (Cluc) on a separate transcript, allowing for ratiometric control for transfection variance (Fig. 1b). We tested this RADARS design along with a scrambled guide control, in the presence or absence of exogenous ADAR2 deaminase domain with hyperactive mutation E488Q and specificity mutation T490A fused to the MS2 coat protein (MCP-ADAR2dd(E488Q, T490A))^17,24^. RADARS-expressing plasmids were co-transfected into HEK293FT cells with either a EGFP expressing plasmid or a control plasmid. We observed that the RADARS sensor resulted in up to a 5-fold increase in the normalized luciferase value, when only relying on endogenous ADAR, and a 51-fold activation in signal (fold change of luciferase expression in the presence of target/in the absence of target, see Methods) when supplemented with exogenous MCP-ADAR2dd (E488Q, T490A) (Fig. 1c, Ext. Data Fig. 1a). To confirm that payload expression is dependent on RNA editing, we harvested RNA from cells and quantified editing with next generation sequencing. We observed a ∼ 24-fold increase in the editing of the UAG stop codon in EGFP targeting sensor, but negligible increase in the editing of a non-targeting sensor (Ext. Data Fig. 1b).

To enable single-cell measurements, we next engineered a RADARS design with a fluorescent payload (Fig. 1d). We designed the fluorescent RADAR sensor as a single transcript, containing mCherry as normalization control, a guide targeting the EGFP transcript, a self-cleaving peptide sequence T2A, and a payload mNeon protein. We transfected HEK293FT cells with combinations of RADARS, exogenous MCP-ADAR2dd (E488Q, T490A), and a frame-shifted non-fluorescent EGFP target transcript. We quantified fluorescence signals by calculating the average mNeon signal per cell across mCherry-positive cells (see Methods) and observed target-specific induction of mNeon by 21.4-fold (Fig. 1e-f) in the presence of exogenous MCP-ADAR2dd (E488Q, T490A) (p<10^−6^, unpaired t-test). We observed weak but significant activation using endogenous ADAR (p<1e^-5^, unpaired t-test) (Ext. Data Fig. 1d), and no background activation with MCP-ADAR2dd (E488Q, T490A), in the absence of target. To confirm that payload expression is dependent on sensor editing by ADAR, we harvested RADARS RNA from cells and quantified editing via next generation sequencing (Fig. 1g). In the presence of exogenous ADAR, we observed 9.4% editing of the UAG stop codon in the presence of the target, and 0.2% editing in the absence, suggesting that target driven editing drives fluorescent payload expression.

### RADARS sensor optimization

During the validation of RADARS we observed that, for some guides, activation can happen in the presence of exogenous ADAR despite the absence of a target RNA (Ext. Data Fig. 1a). To optimize RADARS and minimize this background, we selected and tested a panel of different ADAR1 and ADAR2 mutants in combination with 69 nt guides targeting a frame-shifted EGFP transcript or iRFP transcript (Ext. Data Fig. 2a-d). We screened full-length human ADAR isoforms (ADAR1 p110, ADAR1 p150, and ADAR2) ^25,26^ and their catalytic deaminase domains, along with specific mutants designed to destabilize ADAR-dsRNA interactions to decrease non-specific editing^17,27^. Whereas our initial construct for exogenous ADAR expression, MCP-ADAR2dd (E488Q, T490A), performed the best on the frame-shifted EGFP transcript, several candidates in our screen had comparable activation upon target co-transfection with reduced background (Ext. Data Fig. 2a-d). As guide choice could affect overall sensitivity of the sensor, we screened top ADAR candidates on multiple guide sequences and targets in an orthogonal panel and observed activation above background only in the correctly matched RADARS and target transcripts (Ext. Data Fig. 2e). ADAR1 p150 had highest fold activation on 3 of the 4 targets, driven by a generally low overall background signal in the absence of target. MCP-ADAR2dd (E488Q, T490A) performed best on the EGFP target, due to its generally high level of absolute signal, but suffered from higher background on the other targets that reduced its overall activation (Fig. 2a and Ext. Data Fig. 2f-i, 3a). Interestingly, ADAR1 p110, while not the best, still had good sensor activation on multiple targets, which is advantageous given the ubiquitous expression of ADAR1 p110 in human tissues.

**Figure 2:**
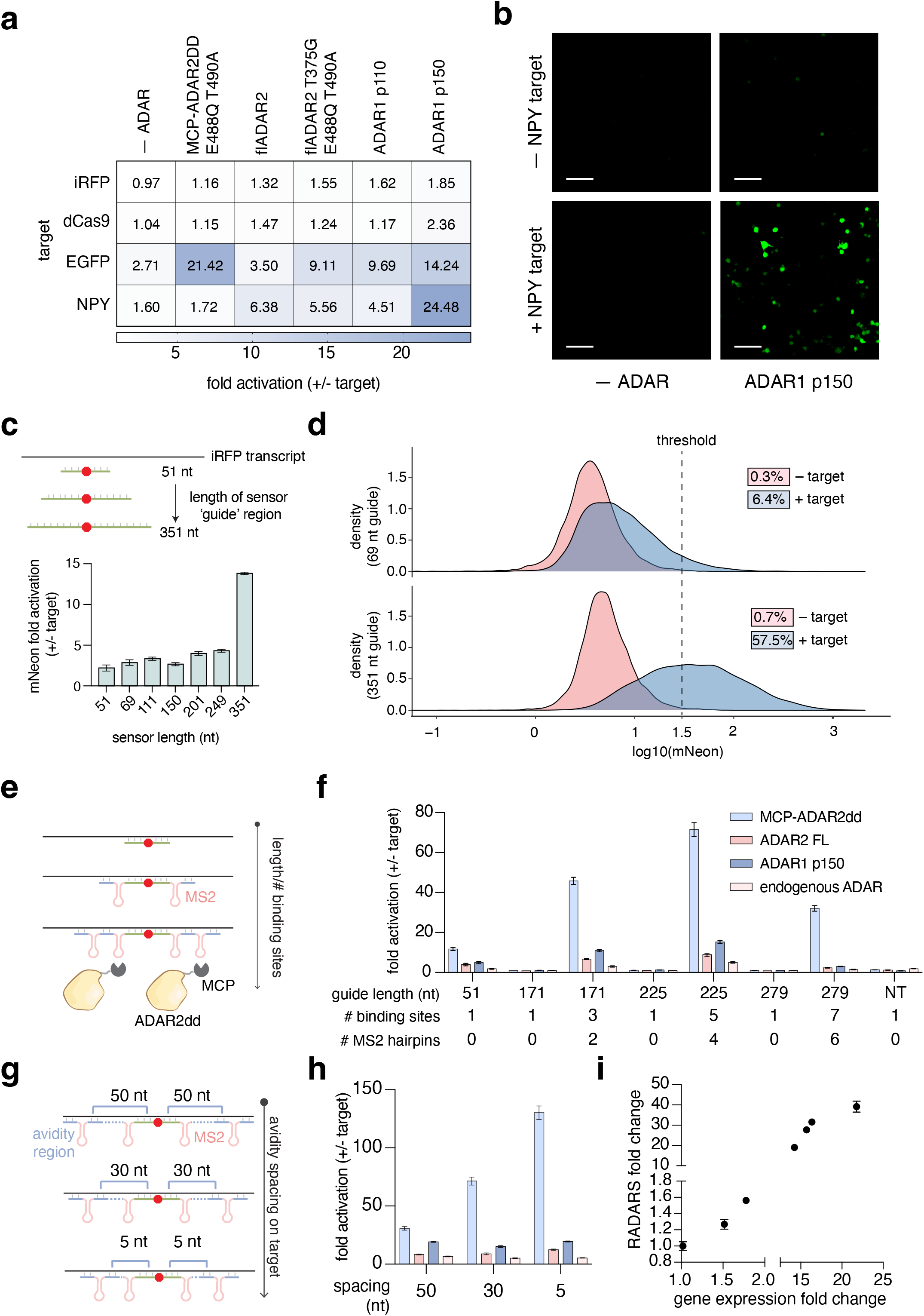
RADARS sensor engineering improves dynamic range and activation. a) Top performing ADAR variants are screened against four different targets in combination with respective RNA sensors. The numbers in the heat map represent ratio fold change. b) Representative images are shown for data in (a) for the neuropeptide (NPY) sensor +/– the NPY target with ADAR1 p150. c) Top: schematic showing different length sensors screened against an iRFP target transcript. Bottom: Bar graph showing increasing sensor activation with increasing sensor length. Sensor activation indicates that normalized fluorescence (mNeon/mCherry) values in the presence of target are divided by normalized fluorescence (mNeon/mCherry) values in the absence of target for each sensor. d) Single-cell image analysis analogous to fluorescence cytometry for iRFP targeting RADARS of guide lengths 69 and 351 nt. Histograms show population density of mNeon expression across all cells for +iRFP (blue) and – iRFP (pink) target conditions. The dotted line shows a constant intensity threshold across all conditions gating individual cells as mNeon(+) or mNeon(–). Colored boxes show % mNeon positive cells for +iRFP (blue) and -iRFP (pink) target conditions. e) Schematics of utilizing MS2 hairpin loops to recruit MCP-ADAR2dd for engineered recruitment of ADAR and increasing binding sites for enhanced hybridization of sensor with target. f) Comparison between sensors containing uninterrupted guides and avidity binding guides containing multiple binding sites separated by MS2 hairpin loops. All guides target the human IL6 transcript with and utilized either endogenous ADAR in HEK293FT cells, exogenous supplemented ADAR1 p150 isoform, full length ADAR2, or MCP-ADAR2dd (E488Q, T490A). Fold change is calculated by the normalized luciferase values (Gluc/Cluc) of the + target condition over – target condition. Data are mean of technical replicates (n= 3) ± s.e.m. g) Schematics of varying avidity spacing on the target in a five site avidity binding sensor. h) Comparison of sensor fold activation between guides of different linker lengths (5 nt, 30 nt and 50 nt) between the binding sites of a five avidity sensor against IL6 transcripts. Data are mean of technical replicates (n= 3) ± s.e.m. i) A seven site avidity binding guide with dual stop codons targeting IL6 is used to quantify the relative expression of IL6 with broad dynamic range. Target expression range is created through a combination of the transient overexpression of tetracycline inducible IL6 (blue dots) and a stable lentivirally integrated tetracycline-inducible IL6 cassette (black dots) in HEK293FT cells. RADARS fold change relative to the basal condition (0ng/mL tetracycline in the integrated HEK293FT cells) is plotted against the IL6 gene expression change as determined by quantitative polymerase chain reaction (qPCR). Data are mean of technical replicates (n= 3) ± s.e.m.

To improve sensor binding stability and target search time, we tested increasing guide lengths centered around a premature stop codon^19^ against a constitutive iRFP target. Sensor activation improved from 2.2-fold to 13.8-fold as guide length increased from 51 nt to 351 nt (Fig. 2c and Ext. Data Fig. 3b). Additionally, we observed a marked shift in the distribution of mNeon expression levels per cell for all guide lengths when target is present (Fig. 2d and Ext. Data Fig. 3c), with a more substantial mNeon(+) population at longer guide lengths. Meanwhile, the percent mNeon(+) cells in the absence of target consistently remained <5% for all guide lengths. At guide lengths of 351 nt, we observed 57.5% mNeon(+) cells in the presence of target, and <1% in the absence of target, suggesting a robust capability to separate cellular populations based on target mRNA expression.

As longer guides may form long double-stranded RNA (dsRNA) duplexes when bound to their targets in cells, we wanted to evaluate whether RADARS elicits an innate immune response. Upon transfection with 249 nt guide RADARS sensors, we evaluated mRNA induction of MDA5 and interferon-β (IFN-β) by qPCR in presence or absence of target and with or without exogenous ADAR1 p150. We found that there was no upregulation of either MDA5 or IFN-β (Ext. Data Fig. 4a-b), suggesting RADARS does not cause immunogenicity in cells. In addition, we investigated whether RADARS with long guide lengths could activate Dicer induced dsRNA cleavage and lead to knockdown of target transcripts. We evaluated target levels by qPCR across four targets and sensors, finding that there was no significant change in transcript levels (Ext. Data Fig. 4c-d). These data are consistent with previous observations that long guides for RNA-editing are not perturbative^19^.

**Figure 4:**
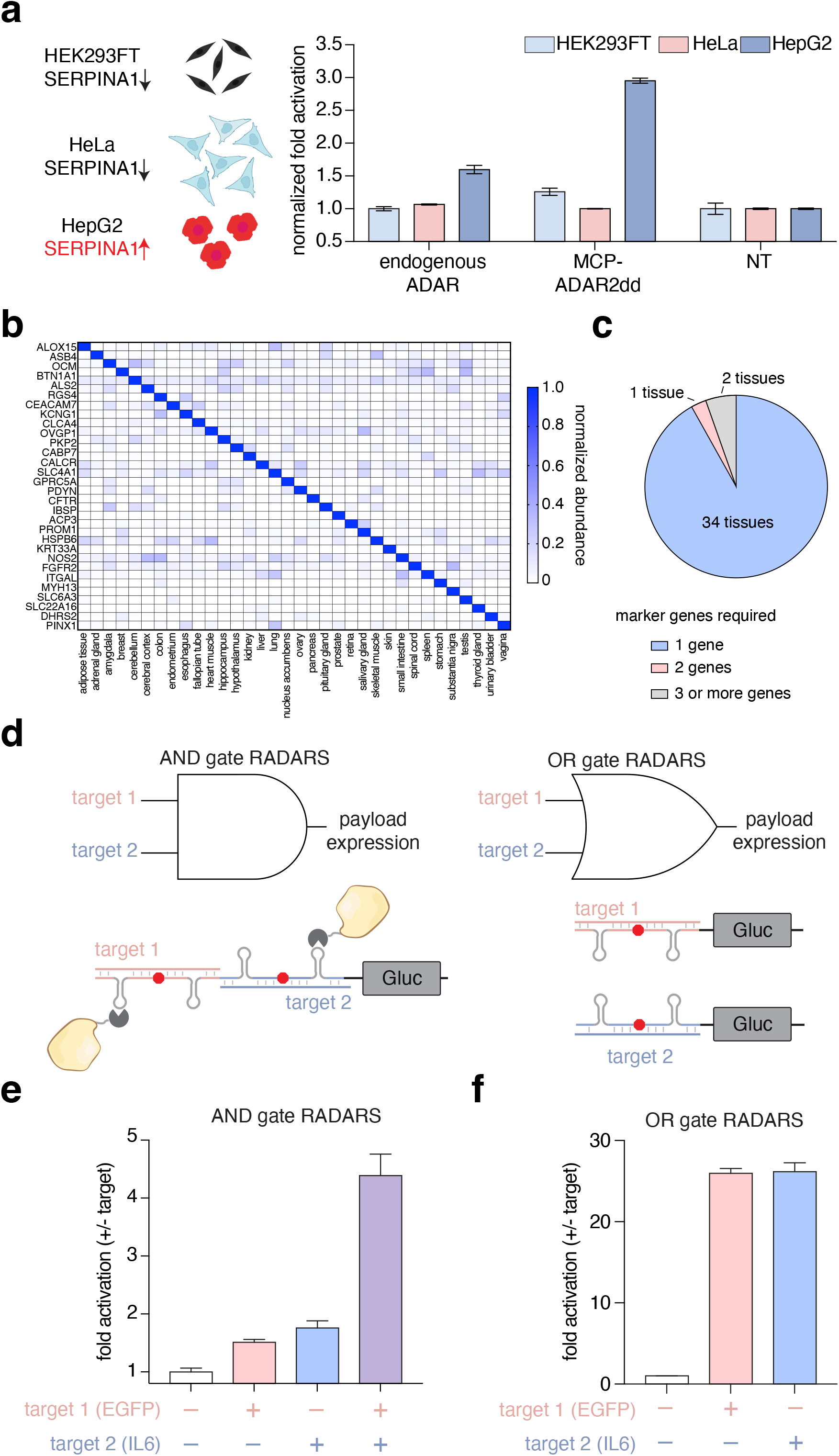
RADARS can be used for cell type classification and complex logic. a) Left: Schematic of SERPINA1 expression difference between HEK293FT, HeLa, and HepG2 cells. Right: The SERPINA1-targeting five site avidity binding sensor is transfected either with or without exogenous MCP-ADAR2dd (E488Q, T490A) in three cell types. Sensor activation is determined by calculating the Gluc/Cluc ratios of SERPINA1 sensor normalized by a scrambled non-targeting guide sensor (NT) to control for protein production/secretion and background ADAR activity differences between cell types, followed by normalization to the Gluc/Cluc ratio in HEK293FT cells. Data are mean of technical replicates (n= 3) ± s.e.m. b) Relative transcript abundance of 34 mRNAs that uniquely define a tissue across 34 different tissue types. c) Summary of the number of genes needed to classify one tissue from the rest. d) Schematic of two input AND gate and OR gate forms of RADARS. e) Normalized sensor activation of AND gate RADARS for EGFP and IL6 transcript inputs across all four possible target combinations. Data are mean of technical replicates (n= 3) ± s.e.m. f) Normalized sensor activation of OR gate RADARS for EGFP and IL6 transcript inputs across different target combinations. Data are mean of technical replicates (n= 3) ± s.e.m.

### RADARS sensor optimization through guide engineering

The two best performing ADARS from our protein screen, ADAR1 p150 and MCP-ADAR2dd (E488Q,T490A), achieved optimal signal through either reducing background or increasing activation. Because MCP-ADAR2dd (E488Q, T490A) had the highest activation with a luciferase sensor (Ext. Data Fig. 5a), we hypothesized that guide engineering strategies to lower background, when coupled with MCP-ADAR2dd (E488Q, T490A), would result in optimal sensor activation. We designed a new sensor targeting IL6 mRNA, a virtually unexpressed transcript in HEK293FT cells^28^.

**Figure 5:**
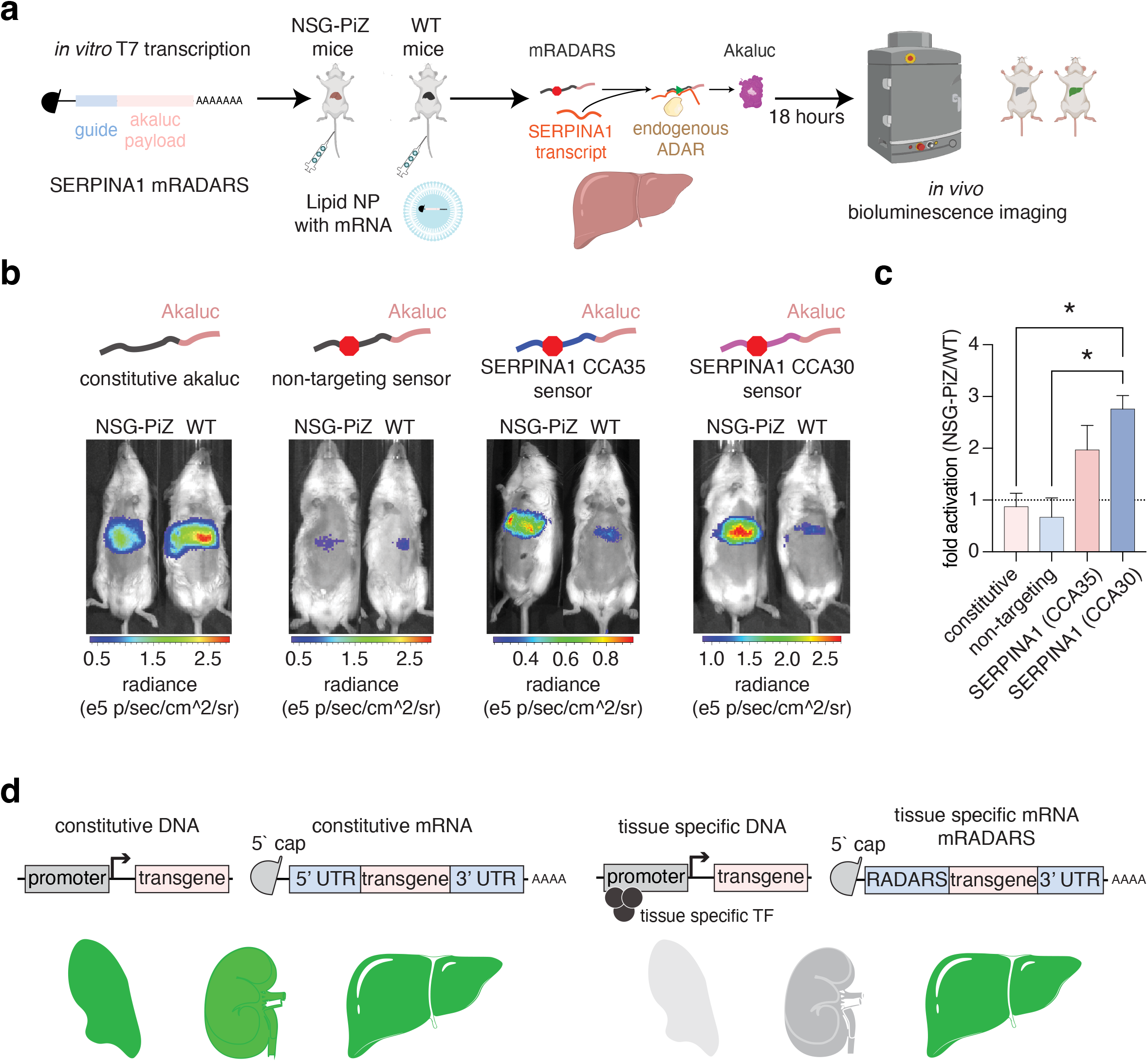
RADARS can be used for synthetic mRNA control *in vivo*. a) Schematic of *in vivo* sensing of the human *SERPINA1* transcript using SERPINA1 mRADARS construct with no exogenous ADAR supplemented. A SERPINA1-targeting sensor mRNA with an Akaluc payload is *in vitro* transcribed with 25% 5-methylcytosine and 0% pseudouridine. Constitutive Akaluc (no stop codon) and a non-targeting guide (with stop codon) sensor constructs are synthesized with the same protocol. All mRNAs are packaged into lipid nanoparticles and tail-vein injected into either wild-type mice or NSG-Piz mice with human SERPINA1 PiZ mutant cassette. *In vivo* sensor activation is measured 18 hours post injection. b) Bioluminescence images of sensor activation for various synthetic mRNA RADARS constructs without exogenous ADAR. c) Akaluc-generated radiance is calculated for the liver and compared between the wild type and the NSG-PiZ mutant mice. The fold change between the NGS-PiZ mice and the WT mice is calculated for each RADARS construct. Significance is determined via a two tailed t-test, N = 2 mice. *, p<0.05. d) Schematic of applying RADARS for control of synthetic mRNA expression in cell-specific contexts. The use of RADARS in this manner is akin to the use of cell-specific promoters for controlling expression of DNA vectors.

We repeated our investigation of increased guide regions with the luciferase sensors targeting an IL6 transcript and found a significant reduction in the fold-change of RADARS activation past 81 nt guides using the MCP-ADAR2dd (E488Q, T490A) construct (Fig. 2e-f). This reduction was due to an increase in background signal in the absence of target, potentially due to readthrough of the stop codon within longer guide regions (Ext. Data Fig. 5b-e). To test this hypothesis, we engineered the guide region to block aberrant translation by introduction of MS2 hairpin loops^29^, which provides the added benefit of recruiting the MCP-ADAR2dd(E488Q, T490A) protein to the guide:target duplex (Fig. 2e-f).

We evaluated these engineered guides with multiple MS2-interspersed binding sites, termed avidity binding guides (10 nt spacing on the target), by varying the number of MS2 loops and binding sites on the guide and comparing to designs with the uninterrupted guides of equal length (Fig. 2e-f). RADARS activation was highest with five site avidity binding guides, achieving ∼70 fold activation and substantially lower background than the uninterrupted guide designs (Extended Data fig. 5f-g). To further explore the avidity binding guide concept, we varied the spacing between the binding sites (5, 30 and 50 nt), finding that binding sites closer together on the target transcripts resulted in the highest activation using the five site avidity RADARS design (Fig. 2g-h). We also explored whether the avidity binding guide improvements could be combined with additional stop codons to further reduce background, and found that introduction of a second UAG stop codon in the guide targeting another CCA site within the guide:target duplex (Ext. Data Fig. 5h) increased the fold activation for the seven site avidity binding guide (Ext. Data Fig. 5i-j). This improvement was driven both by decreased background activation and increased target-specific activation of payload (Ext. Data Fig. 5j-k). Lastly, we explored the mismatch tolerance of RADARS to investigate targeting flexibility. We designed 16 targets, covering all nucleotide changes to either the 5` cytosine or the 3` adenosine (NCN) and found that with the exception of the ACA, ACU, and ACG targets, the five site avidity binding guide RADARS design had optimal sensor activation across all the target motifs (Ext. Data Fig. 6a), expanding the RADARS targeting range.

Avidity binding guides improved performance for all exogenous ADAR constructs, but showed maximal performance increases with MCP-ADAR2dd(E488Q, T490A) supplementation. However, avidity binding guides with five or seven binding sites could produce detectable activation in the presence of only endogenous ADAR with up to 7-fold activation (Fig. 2f), suggesting the recruitment of endogenous ADAR is sufficient for sensor function, which is advantageous given the ubiquitous expression of ADAR1.

### Quantitative detection with RADARS across a high dynamic range

After optimizing the RADARS design, we sought to explore the quantitative response of the system. We produced a wide range of expression levels with both transfected and virally integrated versions of our inducible IL-6 expression system and measured the luciferase response of the seven site avidity binding guide with dual stop codons. We found that the RADARS luciferase activation was linearly correlated with the concentration of the target transgene as confirmed by qPCR (Fig. 2i and Ext. Data Fig. 6b, R^2^ = .96, p<0.001). Correspondingly, RNA editing of the first stop codon in the RADARS guide had a strong correlation with the gene expression level (Ext. Data Fig. 6c), showing RADARS can quantitatively sense transcripts at both the RNA editing and payload level.

### Tracking transcriptional states with RADARS

We next applied RADARS to track transcriptional changes. First, we applied RADARs to sense transcriptional downregulation via siRNA. We targeted *NEFM* and *PPIB* with siRNA constructs in HEK293FT cells (Fig. 3a), followed 24 hours later by transfection with fluorescent RADARs sensors. We validated that siRNA constructs specifically and significantly reduced transcript levels via qPCR and found that RADARS enabled robust detection of the corresponding transcript knockdown (Fig. 3b). While RADARs had slightly reduced signal as compared to qPCR, both RADARs sensors had significantly reduced signal when transcript expression was knocked down (p<0.05, unpaired t-test), with the relative order of gene expression change preserved between *PPIB* and *NEFM* sensors. These results suggest that RADARS is sensitive to endogenous transcript downregulation, with the power to detect <50% changes in endogenous gene levels.

**Figure 3:**
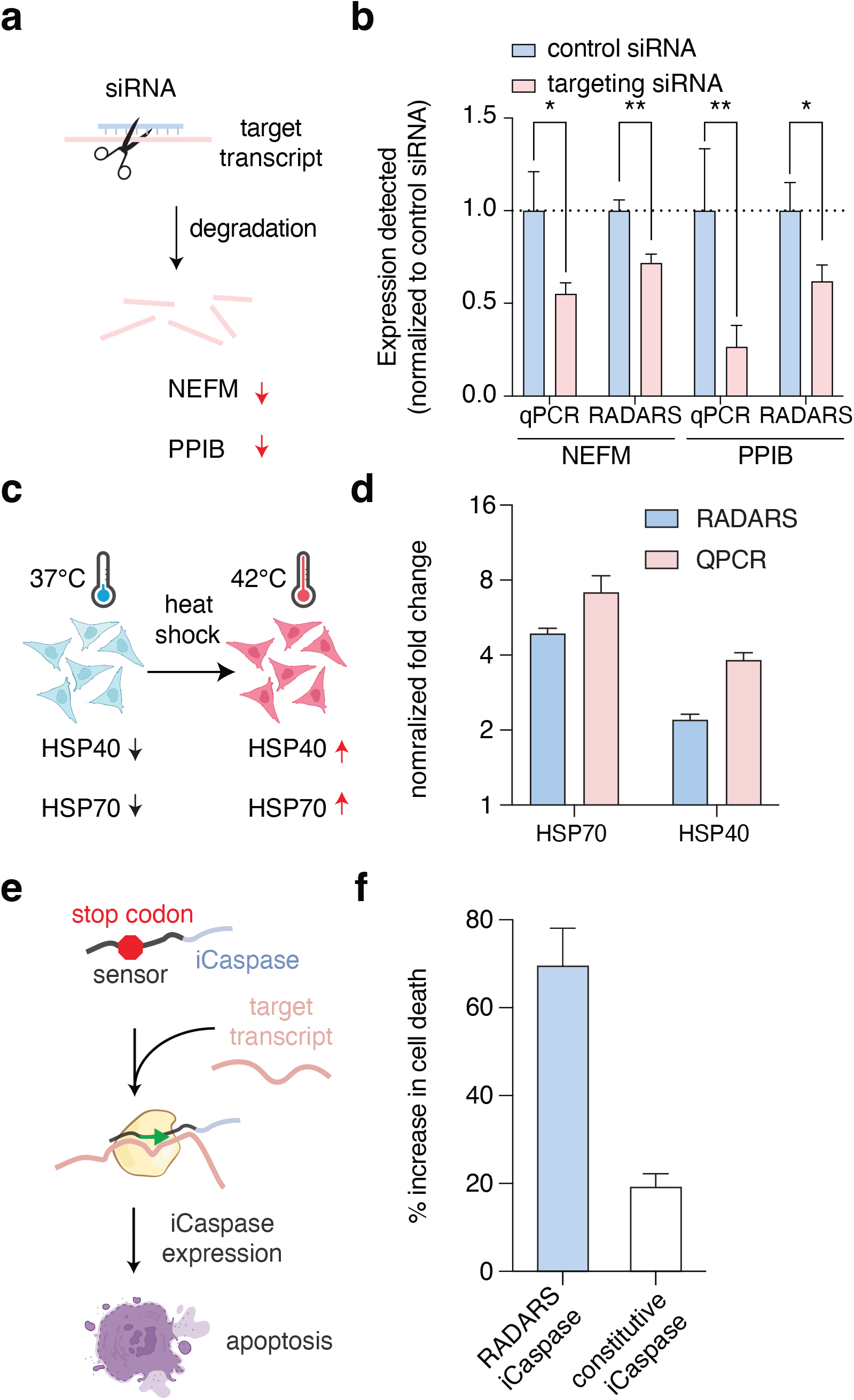
Application of RADARS to tracking endogenous transcript levels. a) Schematic of siRNA knockdown of endogenous transcripts. b) qPCR and fluorescent RADARS detected expression difference between siRNA targeting PPIB or NEFM versus a control non-targeting siRNA in HEK293FT cells. For RADARs sensor activation is calculated for the targeting siRNA and normalized to the control siRNA. Data are mean of technical replicates (n≥ 3) ± s.d. c) Schematic of upregulation of heat shock protein family gene (HSP40 and HSP70) during heat shock at 42 degrees Celsius. d) qPCR and RADARS detected expression differences of HSP40 and HSP70 between the 37 degrees Celsius (control) and 42 degrees Celsius (heat shock) groups. HSP70 and HSP40 targeting sensors, either alone or along with MCP-ADAR2dd (E488Q, T490A), are transfected into HeLa cells followed by 24 hours at 42°C or 37°C. Sensor activation is calculated between the 42°C and 37°C groups and is normalized to a sensor with a scrambled non-targeting guide (NT) to control for changes in protein production. Data are mean of technical replicates (n= 3) ± s.e.m. e) Schematic of an IL6-responsive caspase using RADARS with a five site avidity binding guide targeting the human IL6 transcript and a Caspase 9 payload. f) Fold change of cell death in response to RADARS activation by IL6 transcript detection. Positive control sensors involve a scramble guide sequence in front of the iCaspase with no stop codon in frame. Cell death fold change is determined by calculating the fold change of cell viability in the + target compared to the – target conditions. Data are mean of technical replicates (n= 3) ± s.e.m.

Second, we applied RADARs to sense upregulation of heat-shock family genes in response to cellular heat shock. We designed guides targeting the *HSP70* and *HSP40*, two dynamic heat-shock response proteins, and transfected them into HeLa cells, followed by heat-shock exposure (Fig. 3c). We found that RADARS had a 4.9 fold and 2.2 fold activation for detecting *HSP70* and *HSP40*, respectively, compared to a 7.2 fold and 3.8 fold increase in transcript expression level as validated by qPCR (Fig. 3d). In addition, we tested a previously validated HSP70 IRES-based toehold sensor and adapted the payload into luciferase for direct comparison with HSP70 RADARS and we found no significant detection of heat-shock by the IRES sensor (unpaired t-test, Ext. Data Fig 6d)^14^. These results suggest that RADARs is sensitive to upregulation of endogenous genes, and preserves relative gene expression changes.

### RADARS is compatible with other protein payloads

Having demonstrated RADARS delivery of luminescent and fluorescent proteins as target-specific reporters, we next evaluated whether the modular nature of RADARS allows for diverse payloads that might enable other applications. To apply RADARS for cell state-specific killing, we engineered a payload with the therapeutically relevant iCaspase-9^30^ (Fig. 3e). We found that a RADARS with a IL6-targeting dual stop codon 7 site avidity binding guide and a caspase payload selectively killed IL-6 expressing cells, with minimum toxicity in the absence of IL-6 induction (Fig. 3f and Ext. Data Fig. 7a).

The modularity of protein payloads also allows for small payloads, such as HiBiT payload^31^, that allow for *in vivo* circularization of transcripts. Circular RNAs present a platform for enhanced residency time and minimum immunotoxicity^18^ and we hypothesized that RADARS with small payloads could be circularized to take advantage of these properties (Ext. Data Fig. 6e). We expressed a circular RADARS via a dual Twister ribozyme system^32^ with a HiBiT payload and an IRES to allow for protein expression. We found that circular RADARS expressed the HiBiT in a target (IL6) specific manner (Ext. Data Fig. 7b). For signal amplification, we augmented these circular RADARS as endless RADARS (endRADARS) by removing the stop codon after the payload and inserting 2A peptides on either end of the HiBiT tag, allowing for expression via rolling circle translation (RCT)^33^. We found that endRADARS can express protein in a target specific manner with minimum background leakage (Ext. Data Fig. 7c).

### Cell type sensing with RADARS

Cell type differences represent the primary variation of gene expression in tissues. We utilized RADARS to demonstrate separation of different cell types via marker gene expression. First, to identify marker transcripts for cell type distinction, we performed a differential gene analysis between HEK293, HeLa, and HepG2 cells (Ext. Data Fig. 8a), selecting *SERPINA1*, a liver serine protease inhibitor with therapeutically relevant pathogenic variants ^34^, as a marker only expressed in HepG2 cells and not the other cell lines. We designed a panel of RADARS with guides targeting *SERPINA1* and tested their ability to distinguish between HepG2 and HeLa cells, finding that the CCA30 guide design had the highest activation fold-change between HepG2 and HeLa cells (Ext. Data Fig. 8b). We transfected the *SERPINA1* (CCA30) targeting sensor into all three cell types alongside a non-targeting scrambled sensor designed to control for background ADAR editing, transfection variance, protein production, and secretion differences between the three cell types. We found that the *SERPINA1* targeting sensor could distinguish HepG2 from HEK293FT and HeLa cells with either exogenous MCP-ADAR2dd (E488Q, T490A) or endogenous ADAR (Fig. 4a). Corresponding RNA editing data on the *SERPINA1* sensors and non-target sensors also showed a similar trend, corroborating the specific activation of luciferase in HepG2 cells (Ext. Data Fig. 8c).

While detection of a single marker transcript is useful for distinguishing many cell types, we sought to analyze if more transcripts might be necessary for differentiating all potentially tissue types. Analysis of public tissue gene expression data^35^, shows that 34 out of 37 tissues could be distinguished with a single gene that is 3-fold more expressed over all other tissue types (Fig. 4b), with 3 additional tissues classified by combinations of 2 or 3 marker genes (Fig. 4c). Thus, we investigated if the flexibility of RADARS and orthogonality of guides could enable simple construction of AND/OR logic gates to expand sensing to more than one gene (Fig. 4d). To generate a rudimentary AND gate, we connected two single 51 nt guides, targeting EGFP and IL6 respectively, in tandem with an MS2 hairpin loop. However, this design performed poorly, due to a combination of low signal and background readthrough (Ext. Data Fig. 8d). To improve AND gate signal, we replaced each single binding site guide with a five site avidity binding guide, and found that the resulting AND gate sensor behaved in a target specific manner, requiring both targets to reach full activation (Fig. 4e), with only minor leakage in the single-target conditions. To engineer OR gate logic, we co-transfected two five-binding site avidity sensors targeting EGFP and IL6 and saw that the sensors respond to EGFP or IL6 target transcripts in a manner consistent with an OR gate (Fig. 4f). In total, these results suggest the modularity of RADARS enables logical computations to be performed on mRNA in living cells, which could allow targeting of protein payloads to molecularly defined cell types.

### *In vivo* delivery of mRNA RADARS

Synthetic mRNAs are a useful therapeutic modality, but there are no methods to control their payload expression in a cell-specific manner. We explored the application of synthetic mRNA RADARS (mRADARS) for cell-specific expression in a mouse model that expresses human *SERPINA1* transcripts in mouse hepatocytes. When delivering mRNA, incorporation of base modifications such as 5’ methylcytosine (5mc) and pseudouridine (Ψ) are essential to reduce host immune responses^36^ (Ext. Data Fig. 9a), but these modifications may interfere with ADAR activity, impacting mRADARS function. Using our inducible IL6 system to measure sensor activation, we assayed a range of 5mc or Ψ ratios during mRADARS in vitro transcription and co-transfected modified IL6 sensing mRADARS with exogenous ADAR, as either a plasmid or mRNA (Ext. Data Fig. 9b-c). We found that increased Ψ amounts reduced RADARS activation, while 5mc was more tolerated with 25% incorporation of 5mc having the highest signal activation. To model liver-specific cell targeting *in vitro*, we expressed the human *SERPINA1* transcript in murine Hepa-1-6 cells, in vitro synthesized the top CCA *SERPINA1* sensors as mRNA with 25% incorporation of 5mc, and transfected mRNA sensor alone into Hepa-1-6 cells. We found that both *SERPINA1* sensors targeting CCA30 and CCA35 are able to recruit endogenous ADAR in Hepa-1-6 cells to sense induction of *SERPINA1* transcripts (Ext. Data Fig. 9d).

To test mRADARS *in vivo* we synthesized the CCA30 and CCA35 guides in a RADARS construct that expresses Akaluciferase (Akaluc), which allows for facile non-invasive luminescent imaging to confirm cell-specific RADARS activation (Fig. 5a)^37^. We delivered these mRADARS sensors into either NSG-PiZ mice, which express a humanized version of *SERPINA1*, or wild-type mice, which only express mouse Serpina1 and do not have binding sites for CCA30 and CC35 sensors. To test if *in vivo* sensing could rely only on endogenous ADAR expression for editing, we did not deliver exogenous ADAR2dd. To control for mRNA delivery and expression differences between mouse strains, we also designed an Akaluc payload with either a constitutive mRADARS, lacking the stop codon, or a scrambled non-targeting guide and transfected them to both NSG-PiZ and WT mice. The *SERPINA1*-sensing mRADARS designs had significant activation (p = 0.04, N=2 mice, two tailed unpaired t-test) of Akaluc expression in the NSG-PiZ mice relative to the WT mice, and control guides had no significant difference between the two strains (Fig. 5b-c, Ext. Data Fig. 9e). This activation demonstrates that RADARS can be delivered as synthetic mRNA to sense cellular state *in vivo* with endogenous ADAR.

## Discussion

Here we present RADARS, reprogrammable RNA sensors, for selective expression of a payload protein in the presence of a target RNA. RADARS enable rapid design and numerous flexible applications due to the reprogrammable guide region that can be predictably targeted by base pairing rules and a downstream modular payload for protein translation. We validated the feasibility of RADARS on reporter transcripts and optimized both the protein and guide architecture to produce up to 164-fold activation by engineering hairpins and multiple binding sites into the guide. Moreover, we demonstrate that the RADARS sensors have target-specific activation utilizing endogenous ADAR, facilitating applications where exogenous ADAR might not be desired. We also provide a design guide for avidity sensors (Ext. Data Fig. 10) and an easy to use software program to automatically generate avidity sensors for input target sequences (https://github.com/abugoot-lab/RADARS). We showed RADARS are quantitative across a ∼25-fold range of gene expression, adapted RADARS for boolean logic on multiple inputs, and designed payloads for expression-specific cell killing. We then applied RADARS for identifying different cell types, tracing upregulation of genes in heat shock models and downregulation of genes via siRNA knockdown, and minimally-invasive profiling of gene expression in living animals using RADARS delivered via mRNA.

While we have optimized RADAR activation through progressive engineering of ADAR proteins and guide designs, there remain multiple routes for future sensor improvement. While sensor screening can identify top performing guide locations, further work is needed to thoroughly understand the factors influencing guide performance. In the future, large-scale screening efforts can probe the determinants of sensor editing, including secondary structure and sequence properties that affect readthrough, in the presence and absence of targets, ultimately yielding versatile guide design rules. Here we find that MS2 loops can block readthrough, recruit MCP-ADAR2dd, and potentially stabilize the multivalent guide binding regions and prevent self-folding, as has been seen with similar therapeutic RNA editing systems^20^, and it is likely that additional engineering improvements at both the ADAR protein and guide level could further increase editing and sensor output. For applications in real-time RNA monitoring, more work is needed on tuning the RADARS kinetic properties. As sensor turnover is constrained by mRNA lifetime (both target and sensor) and payload protein lifetime, additional engineering of payload and sensor mRNA lifetime will enable high-resolution temporal studies.

The qualities of the RADARS platform, including high sensitivity, reprogrammability, and modularity, make it ideal for a myriad of applications in fundamental and translational research. As the RADARS mechanism allows for a single target transcript to edit multiple sensor molecules, RADARS has intrinsically high sensitivity to low abundance transcripts. Moreover, the reprogrammability and modularity of RADARS enables Boolean logical gates (AND/OR), which may lead to complex combinatorial logical inputs and outputs in the future, allowing for detection of cell types or states characterized by more than just a single marker transcript. For RADARS payloads, multiple different classes of protein outputs can be used, including fluorescent proteins, luminescent proteins, caspases, and small peptides. Caspase payloads enable selective cell killing, which can have numerous applications for therapeutics, such as tumor cell targeting or elimination of aged or fibrotic cells. Additional payload applications include the delivery of transcriptional activators for further signal amplification or recombinases for marking cell states that leverage numerous existing genetic animal models. Lastly, transcription factor (TF) payloads could be used in models of cell differentiation, or triggering TF cascades.

We anticipate many uses of RADARS in research for tracking cell states *in vivo* based on RNA signature, including neuronal activity in response to behavioral stimuli and immune cell response to pathogens. In these efforts, RADARS could be additionally combined with existing transcriptional tools for conferring cell type specificity, such as cell type specific promoters or enhancers to enable combinatorial targeting of relevant cell populations. For specific applications where exogenous ADAR expression might be undesired, such as clinical scenarios, using RADARS sensors that rely on endogenous ADAR is possible, as we show both *in vitro* and *in vivo* that endogenous ADAR can be recruited. As ADAR1 p110 isoform is functional with RADARS and is widely expressed, we anticipate RADARS can function in most tissues with endogenous ADAR1 recruitment.

Importantly, RADARS allows for translational regulation of synthetic mRNAs used in mRNA biological therapeutics (Fig. 5d). Since mRNAs are delivered post-transcriptionally, it is impossible to utilize existing transcriptional regulation (e.g cell-type specific promoters). Thus, RADARS offers a novel avenue to regulate the expression of these mRNA therapies. Overall, RADARS is a reprogrammable biological sensor platform with many applications for biomedical research, diagnostics, and therapeutics.

## Acknowledgements

We would like to thank P. Reginato, D. Weston, and E. Boyden for MiSeq instrumentation support; and R. Desimone and J. Crittenden for support and helpful discussions. We thank the members of the Chen and Abudayyeh-Gootenberg labs for helpful discussions.

## Authors contributions

O.O.A., J.S.G, and F.C. conceived the study and participated in the design, execution, and analysis of experiments. K.J., J.K., D.C., R.K., and Y.Z. designed and performed the experiments in this study and analyzed the data. N.Z. and L.V. helped with guide engineering strategies. K.J., J.K., D.C., R.K, Y.Z., O.O.A., J.S.G, and F.C. wrote the manuscript with help from all authors. K.J., J.K., D.C., R.K., and Y.Z. contributed equally and have the right to list their name first in their CV.

## Data and materials availability

Sequencing data will be available at Sequence Read Archive. Expression plasmids are available from Addgene under UBMTA; support information and computational tools are available at www.radars.bio.

## Figure legends

**Extended Data Figure 1:**
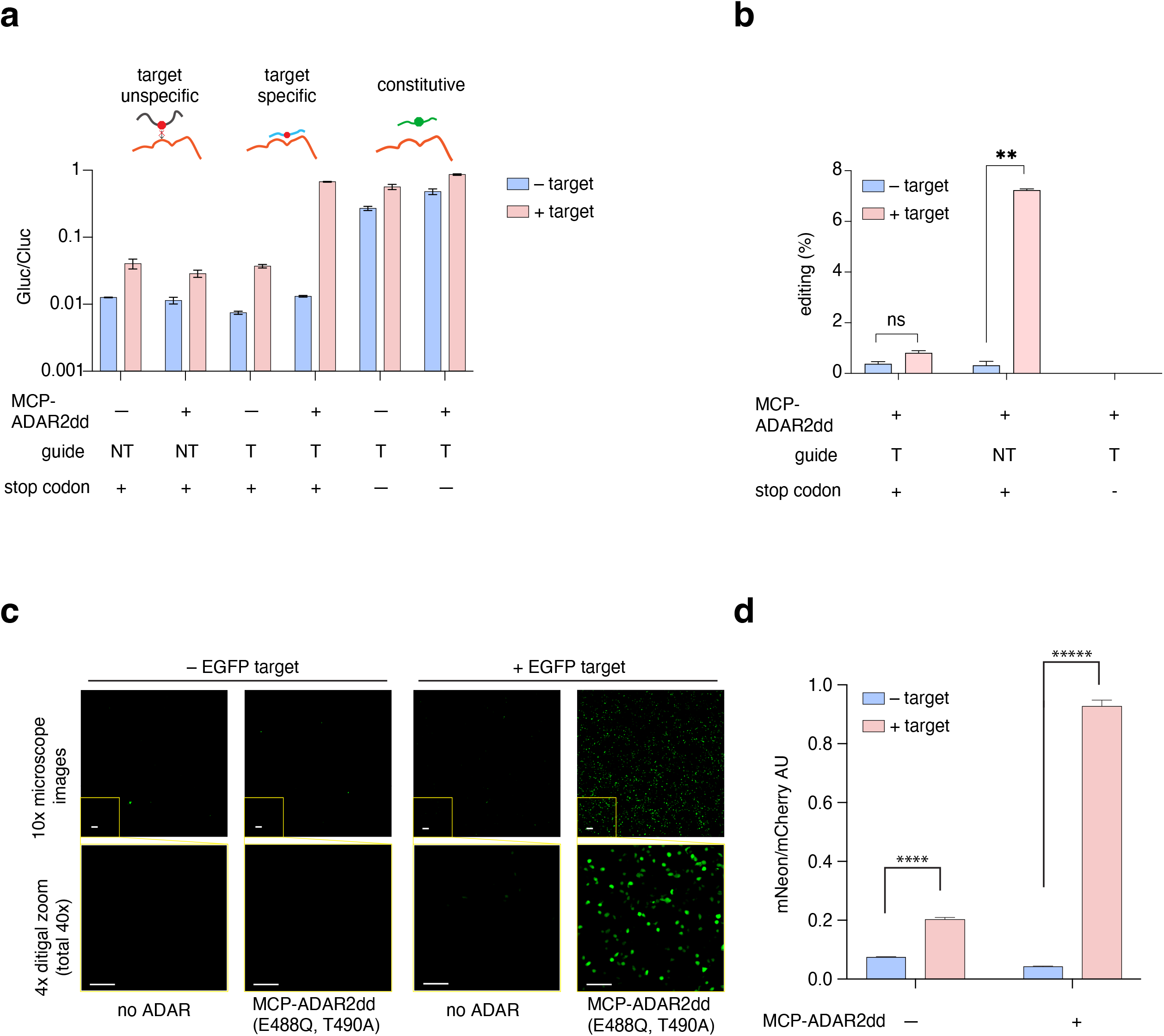
Characterization of luciferase RADARS sensors against EGFP transcripts. a) Comparison of normalized luciferase signal to a non-targeting guide sensor, EGFP targeting sensor, and a constitutive (no stop codon) sensor in the +target and – target groups. b) Corresponding editing percentage of the target adenosine in the UAG stop codon for a non-targeting guide sensor, EGFP targeting sensor, and a constitutive (no stop codon) sensor with exogenous MCP-ADAR2dd (E488Q, T490A). Error bars indicate standard error of the mean. (n=3 technical replicates. ** indicates p<0.01). c) Full 10x images with insets for images shown in Figure 1e. Scale bars are 100 μm. d) Non-normalized fluorescence values (mNeon/mCherry) are shown for all four conditions. Data are mean of technical replicates (n= 3) ± s.d. Significance tests are measured by an unpaired Student’s t-test. ****, p-value <1e-5, *****, p-value <1e-6.

**Extended Data Figure 2:**
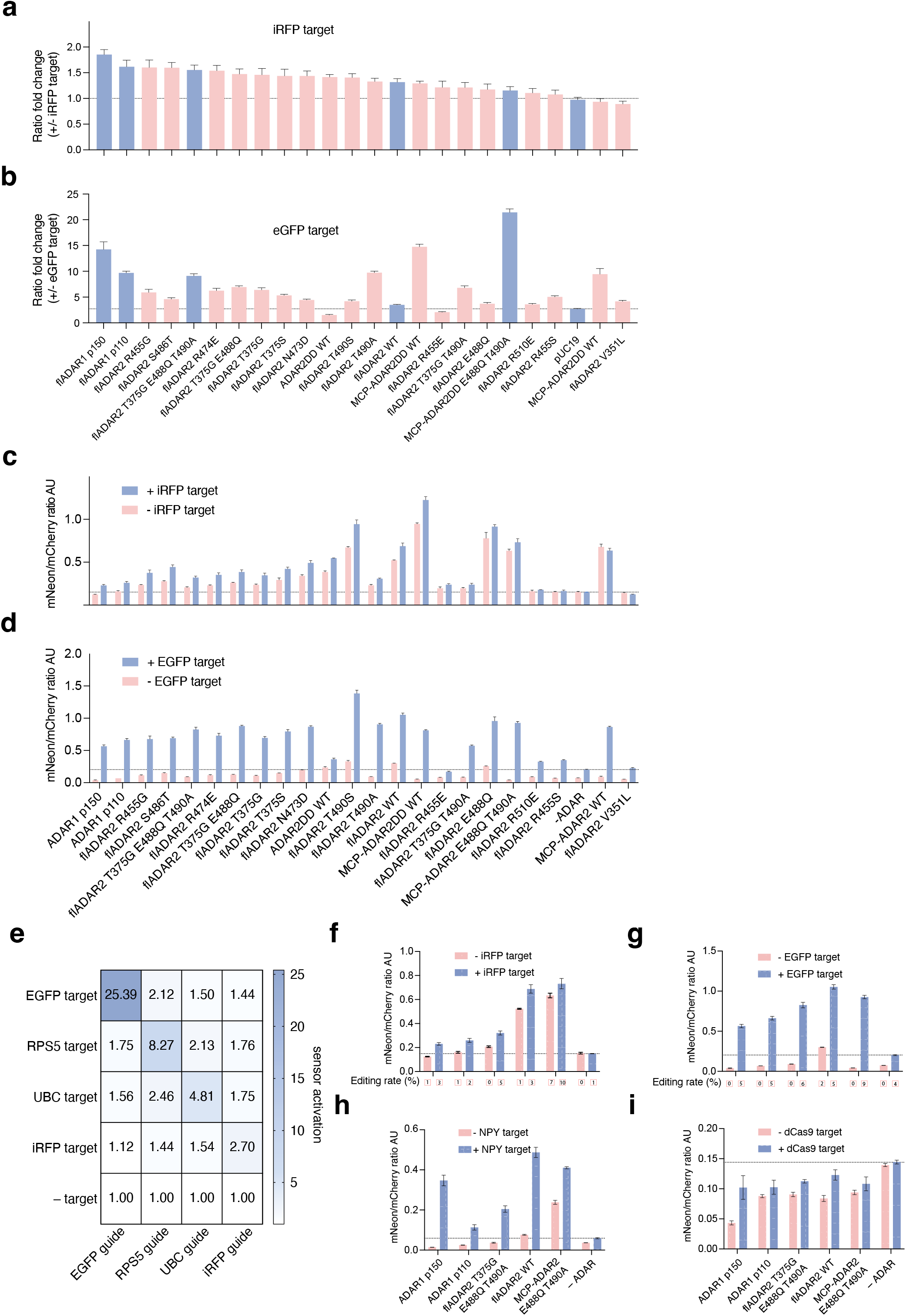
Further characterization of ADAR variants. a) Testing of ADAR variants on a 69 nt iRFP RADARS. Fold change shown indicates fluorescence ratio values (mNeon/mCherry) in the presence of target divided by ratio values in the absence of target. b) Activation of exogenously transfected EGFP targeting RADARS with different ADAR variants and catalytic mutants in the presence of transfected EGFP. Ratio fold change denotes fold change in fluorescence ratio values (mNeon/mCherry, see Methods) between +EGFP and –EGFP target expression. ADAR variants selected for screening across additional targets are shown in blue. c) Non-normalized mNeon/mCherry fluorescence ratio values for data shown in Extended Data Fig. 2a. d) Non-normalized mNeon/mCherry fluorescence ratio values for data shown in Extended Data Fig. 2b. Data are mean of technical replicates (n= 3) ± s.d. e) HEK293FT cells were transfected with ADAR p150 and with plasmids expressing target transcripts and target-sensing RADARS constructs in the combinations shown on the y- and x-axes, respectively. Data shown is fold change calculated as the fluorescent ratios (mNeon/mCherry) in the + target divided by – target (pUC19) conditions. Data are mean of technical replicates (n= 3). f) non-normalized mNeon/mCherry fluorescence ratio values for iRFP sensor with different ADAR variants. Data are mean of technical replicates (n= 3) ± s.d. On the bottom panel, next-generation sequencing data of the RNA sensors for the UAG to UIG conversion. % editing indicates % reads that are A—>I edited. g) non-normalized mNeon/mCherry fluorescence ratio values for EGFP sensor with different ADAR variants. Data are mean of technical replicates (n= 3) ± s.d. On the bottom panel, next-generation sequencing data of the RNA sensors for the UAG to UIG conversion. % editing indicates % reads that are A—>I edited. h) non-normalized mNeon/mCherry fluorescence ratio values for NPY sensor with different ADAR variants. Data are mean of technical replicates (n= 3) ± s.d. i) non-normalized mNeon/mCherry fluorescence ratio values for dCas9 sensor with different ADAR variants. Data are mean of technical replicates (n= 3) ± s.d.

**Extended Data Figure 3:**
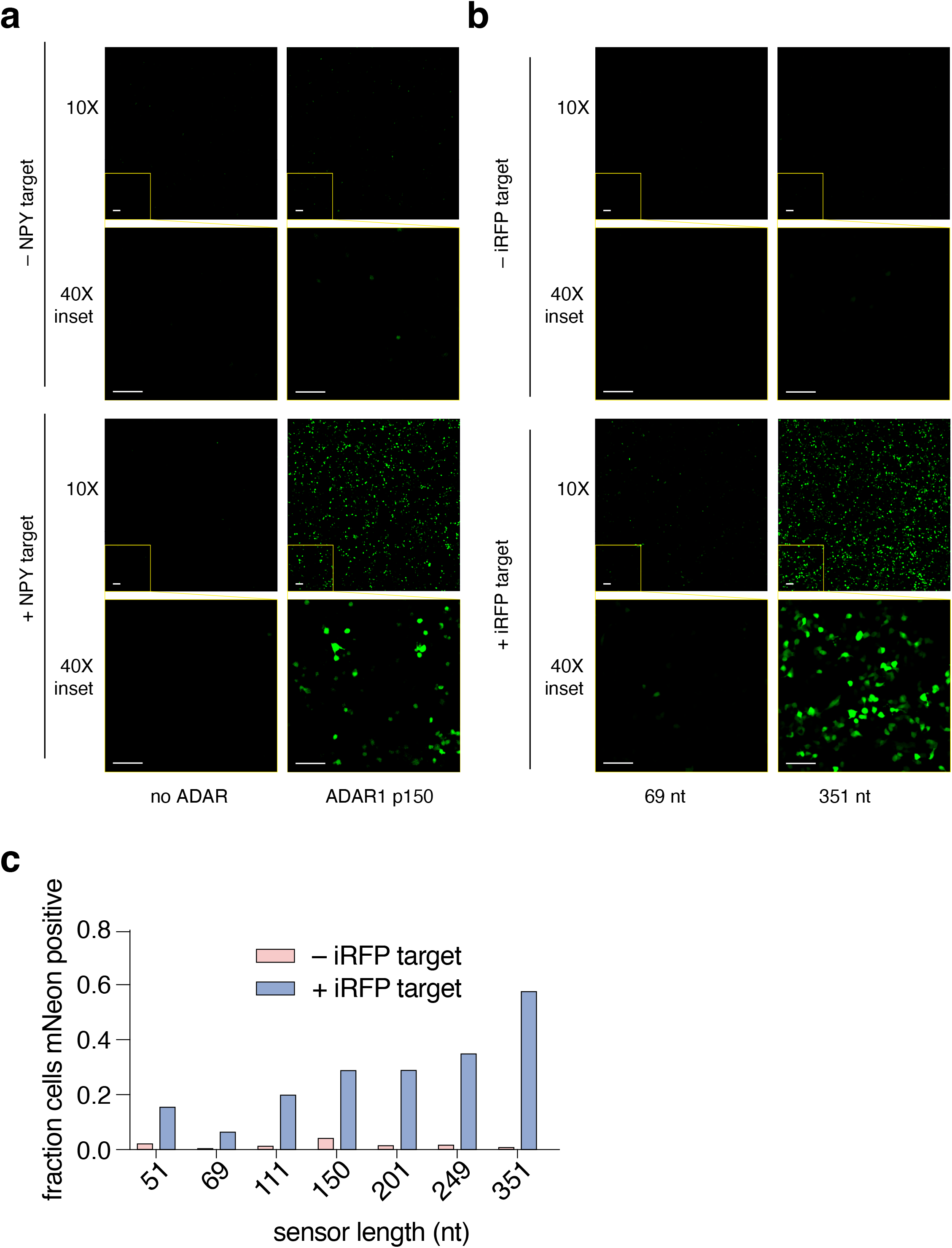
Supporting images of fluorescent RADARS with ADAR1 p150. a) Full 10x images with insets for NPY sensors. Scale bars are 100 μm. b) Full 10x images with insets for iRFP sensors. Scale bars are 100 μm.

**Extended Data Figure 4:**
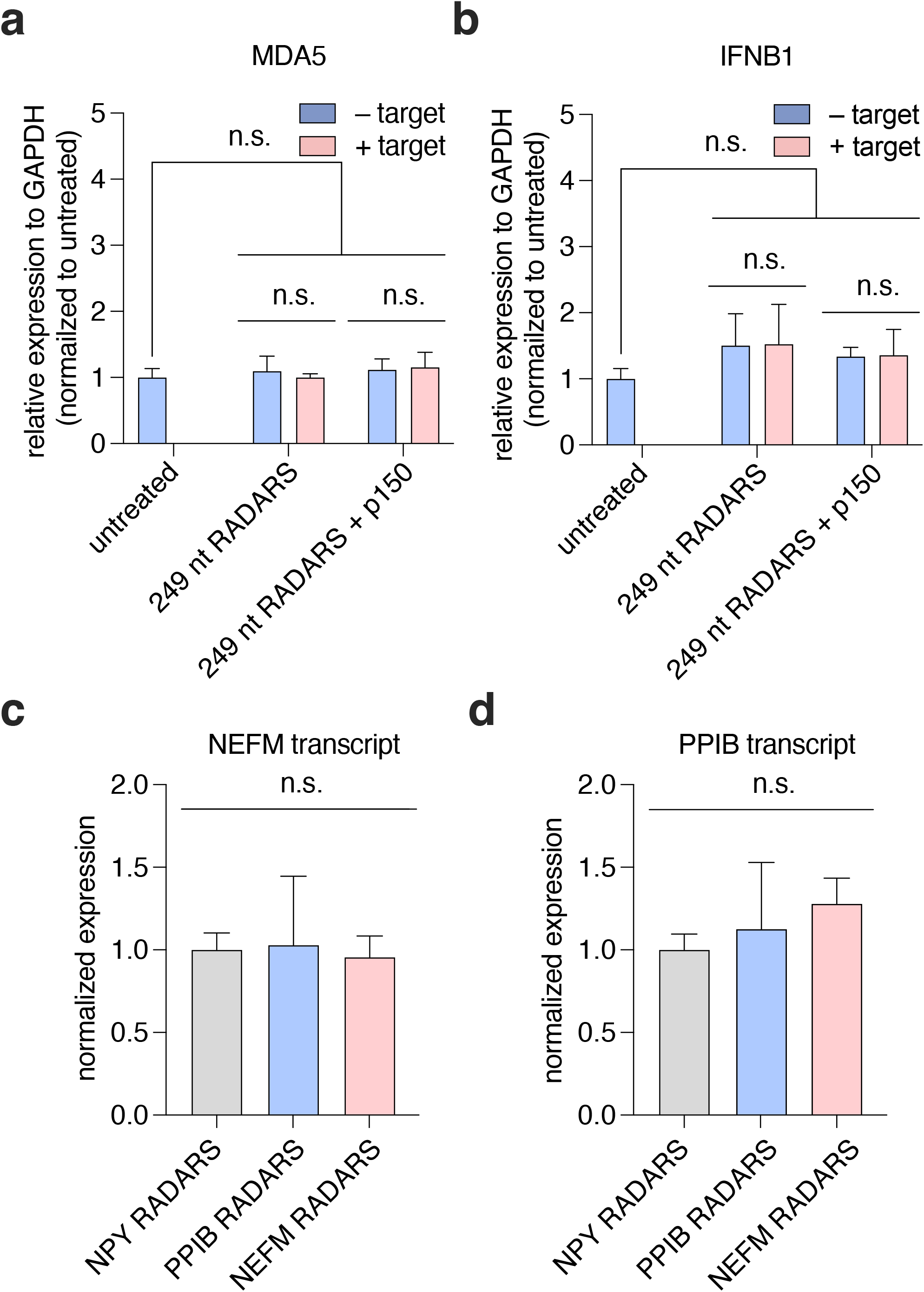
Characterization of RADARS safety regarding immune response and endogenous RNA knockdown. (a and b) Effect of sensor-target duplex formation on the innate antiviral pathways. RADARS sensors were transfected in the presence or absence of complementary target sequences. Total RNA was analyzed using quantitative PCR (qPCR) to determine the relative expression levels of MDA5 (a) and IFN-β (b). (c and d) Effect of sensor-target duplex formation on the abundance of endogenous targeted transcripts. The relative abundance of NEFM and PPIP transcripts upon the transfection of complementary or non-targeting RADARS sensors were assessed by qPCR. Data are presented as the mean ± s.d. (n = 4); unpaired two-sided Student’s t-test, ns, p > 0.05.

**Extended Data Figure 5:**
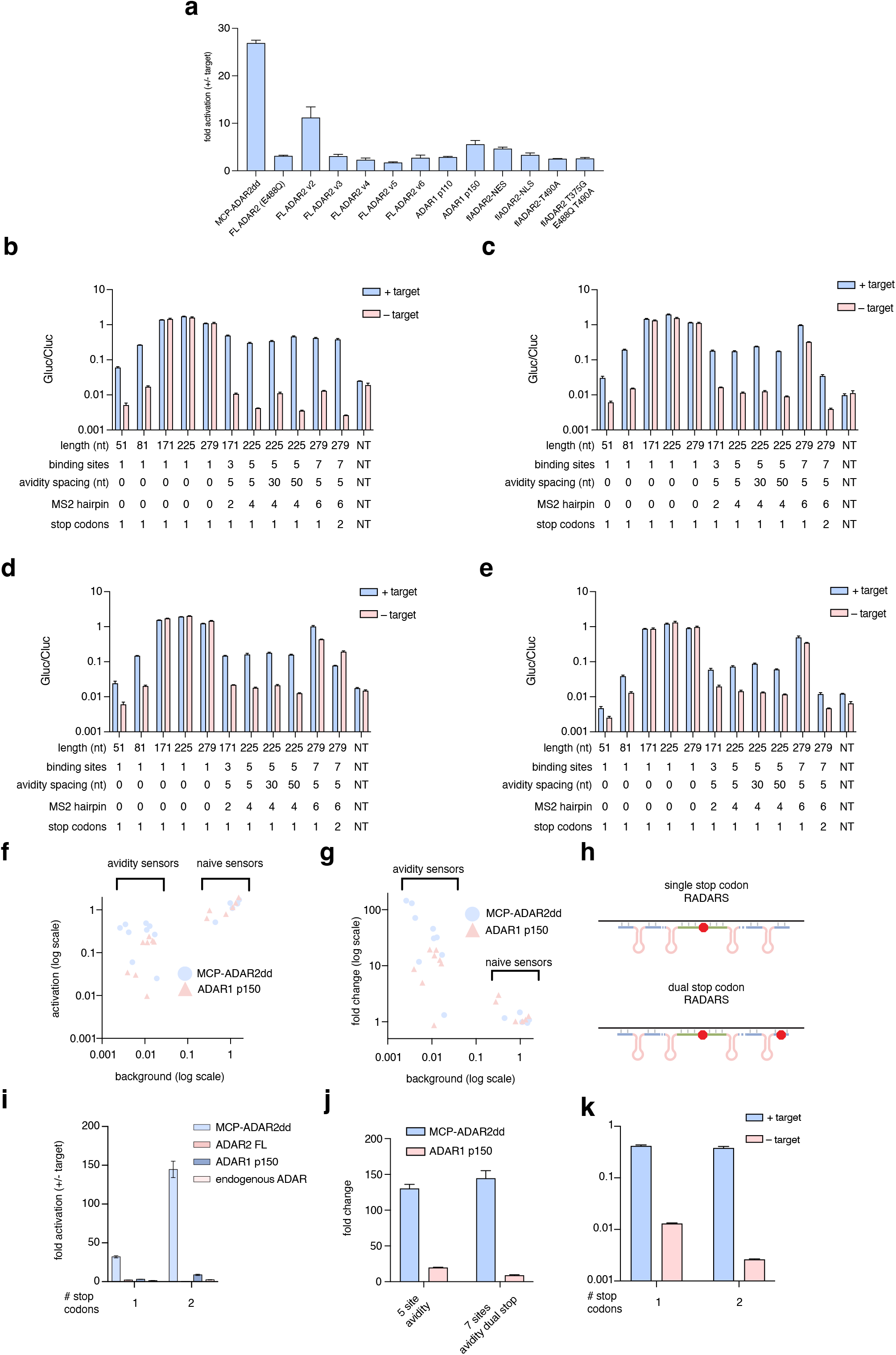
Further characterization of avidity sensors. a) Comparison of the fold change between different ADAR variants on three site avidity binding IL6 sensors. (b, c, d, and e)Normalized luciferase values of panel of sensors in the +target group and –target group for b) MCP-ADAR2dd (E488Q, T490A) exogenous supplementation, c) ADAR1 p150 isoform exogenous supplementation, d) ADAR2 exogenous supplementation, and e) no exogenous ADAR supplementation. f) Comparison of background signal versus activation for avidity sensors with naive guide sensors. g) Activation fold change versus background signal for avidity sensors and naive guide sensors. h) Schematic of dual stop codons in the distant avidity binding site of a seven site avidity binding sensor. i) Comparison of sensor fold activation between a seven site avidity binding guide with a single stop codon and a seven site avidity binding guide with dual stop codons. j) Comparison between five site avidity binding sensors versus seven site avidity binding dual stop codon sensors across MCP-ADAR2dd (E488Q, T490A) and ADAR1 p150. k) Comparison between the activation and background signal of seven site avidity binding single stop codon versus seven site avidity binding dual stop codon sensor.

**Extended Data Figure 6:**
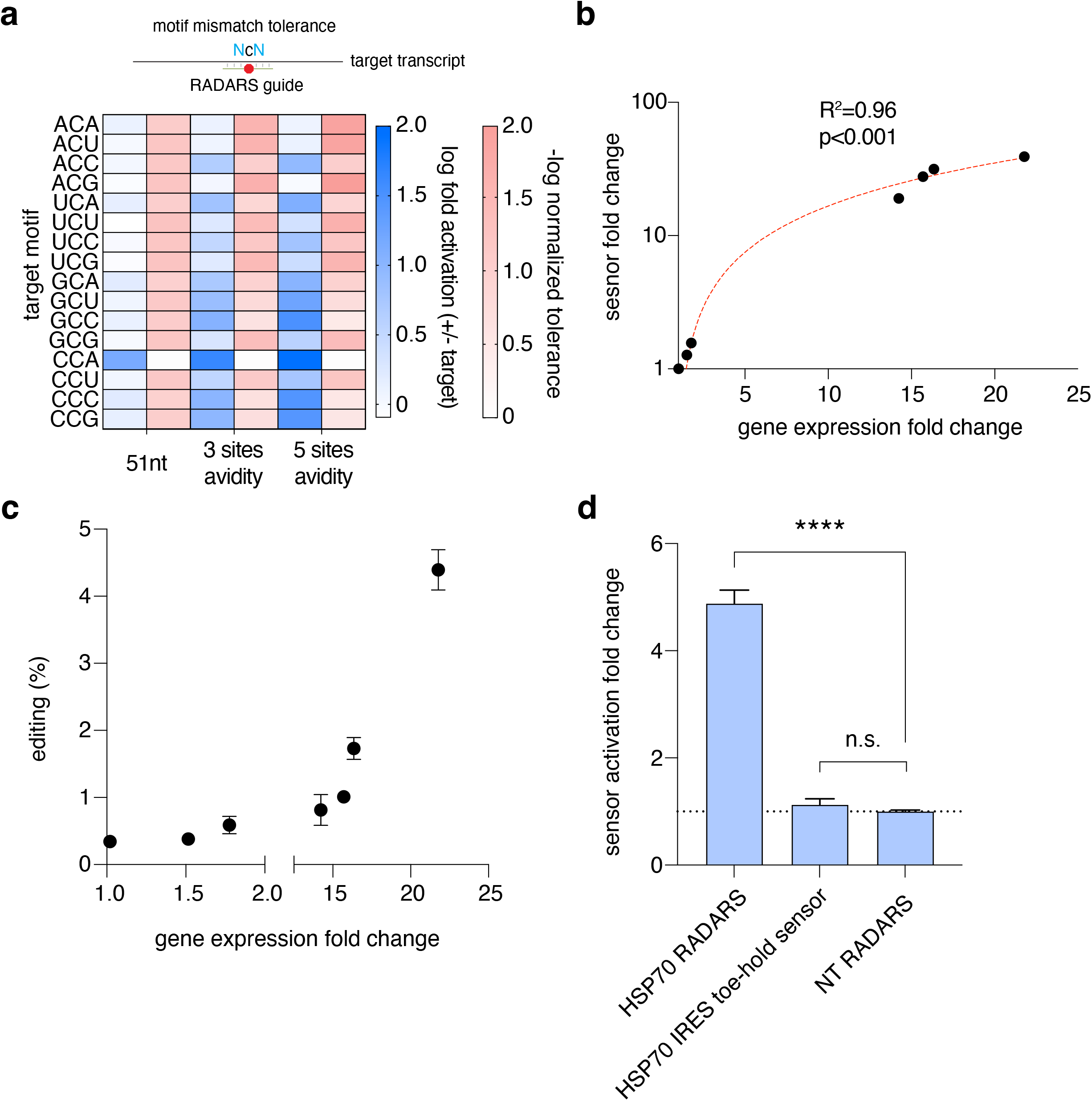
Avidity sensors tolerates target mismatch and enhances activation and dynamic ranges. a) Evaluation of the target mismatch tolerance across all 16 possible target mismatches between the naive 51 nt sensor, three avidity, and five avidity sensor designs. Sixteen possible targets comprise all possible 5` or 3` nucleotide changes from the regular CCA codon (NCN). The left heatmap (blue) shows the log_10_ sensor activation fold change of all three sensor designs across the 16 target mismatches. The right heatmap (red) shows the negative log_10_ normalized tolerance of the different target mismatch relative to the native CCA target. b) A linear regression on the sensor activation fold change against qPCR detected gene expression fold changes. c) Corresponding edit of the adenosine in the UAG stop codon of the sensor across different IL6 gene expression levels. d) Comparison of activation fold changes for RADARS and a previously validated IRES-based toehold sensing constructs^14^ for HSP70 sensing. Data are mean of technical replicates (n= 3) ± s.e.m. Significance tests are measured by a two-tailed Student’s t-test. ****, p-value <1e-4, ns, p-value >0.05.

**Extended Data Figure 7:**
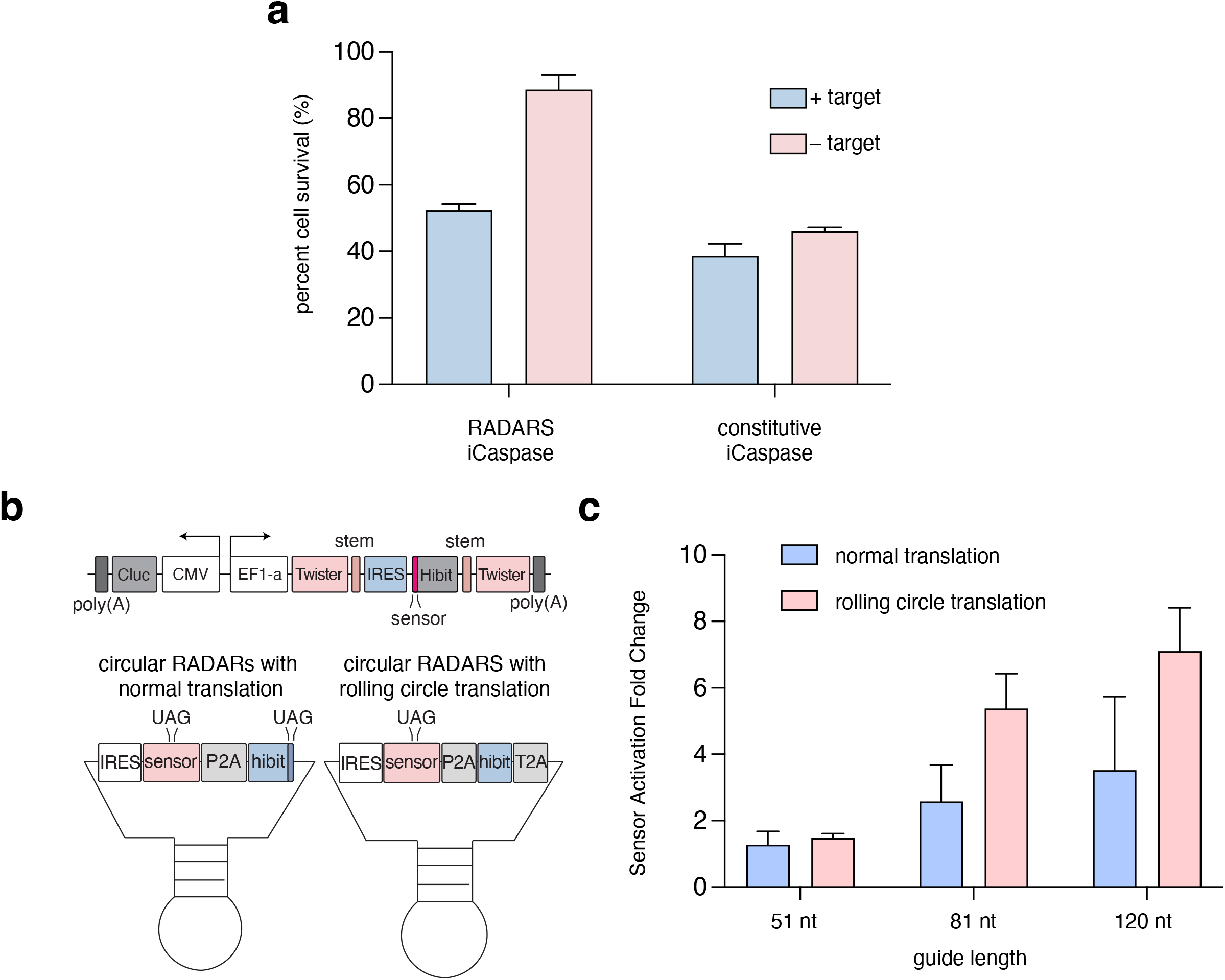
RADARS is compatible with different payloads. a) Comparison of percent cell survival values of IL6 responsive iCaspase RADARS and a no stop codon control in the + target and – target groups. b) Schematic of circular RADARS where RADARS expression is driven by an U6 promoter in a twister ribozyme backbone for intracellular self-circularization. A rolling circle translation version of the circular sensor is engineered by deleting the stop codon at the C-terminal end of the hibit peptide tag and insertion of T2A peptide to allow for ribosomal readthrough in a circular fashion. c) Comparison of guide lengths between 50 nt to 120 nt in a circular RADARS construct for sensor activation fold change upon transgene target (human IL6) induction.

**Extended Data Figure 8:**
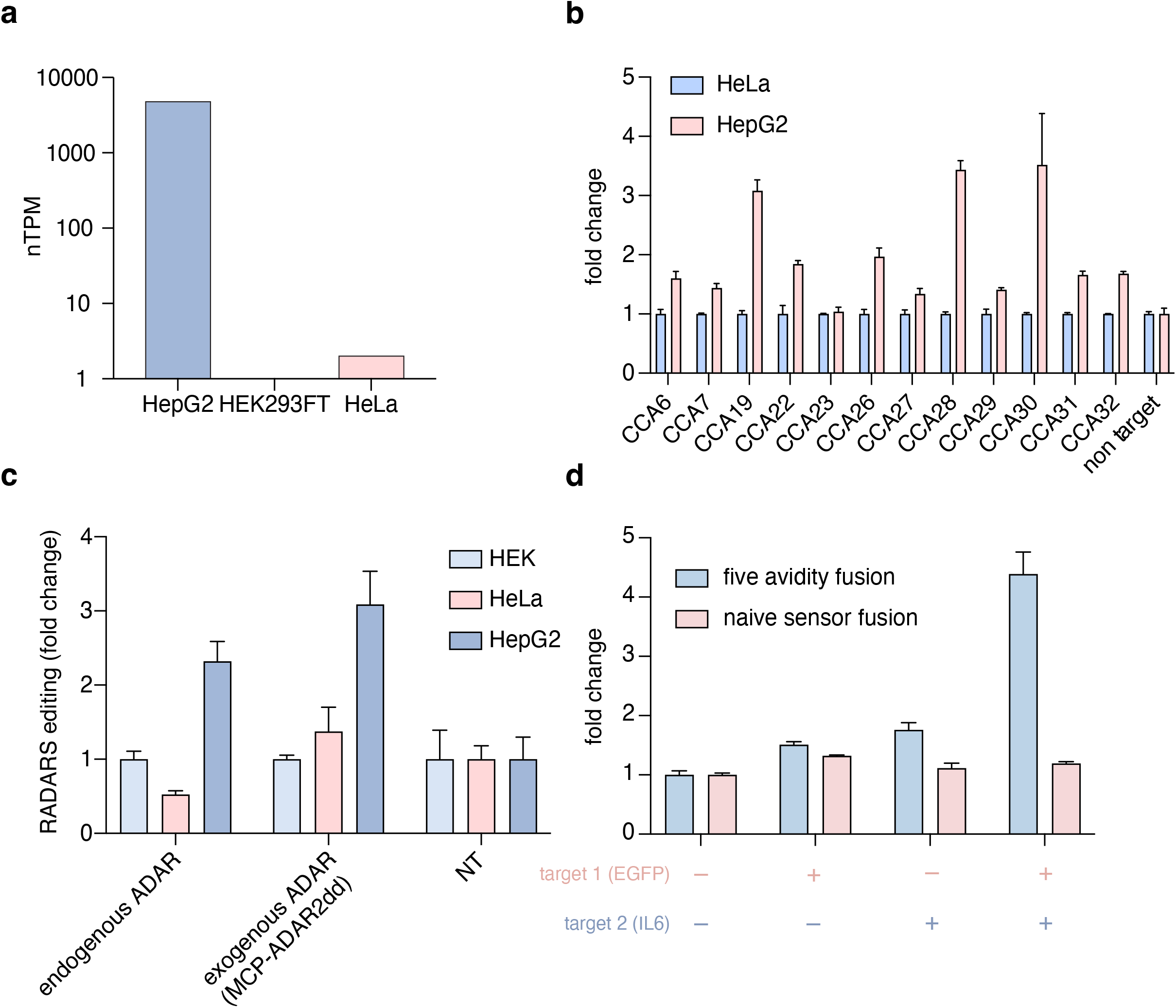
Further characterization of cell typing RADARS. a) Comparison of SERPINA1 expression across HEK293FT, SERPINA1, and HeLa cells. b) Comparison of sensor activation fold changes between HeLa cells and HepG2 cells across SERPINA1 sensors targeting different CCA sites. c) Normalized fold change of editing rate at the adenosine in the UAG stop codon of SERPINA1 5 site avidity binding sensor (CCA30) in different cell types (HEK293FT, HeLa and HepG2). d) Comparison of activation fold change between a continuous 51 nt guide AND gate sensor and 5 site avidity binding guide AND gate sensor across all four combinations of IL6 and EGFP target induction.

**Extended Data Figure 9:**
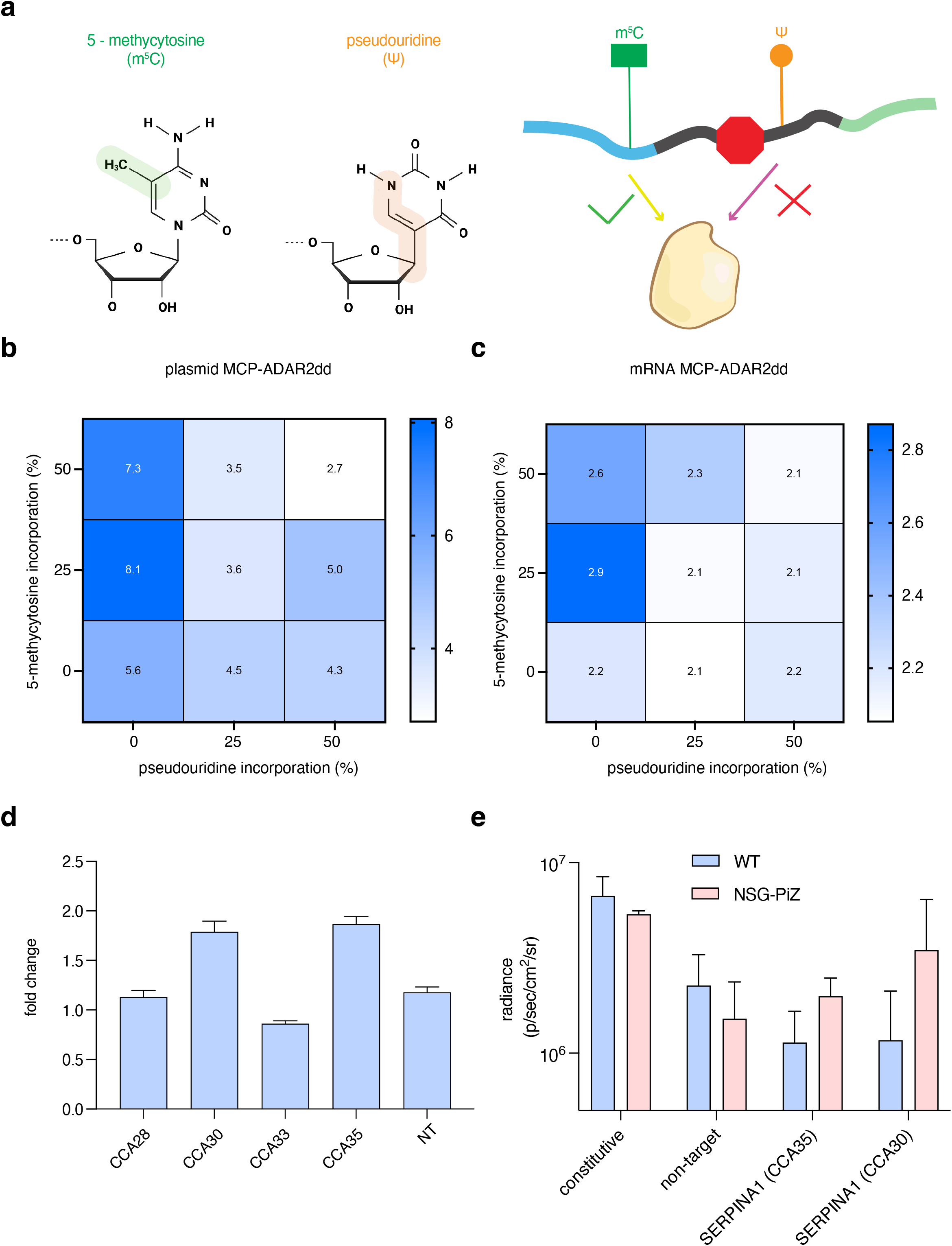
Further characterization of mRNA RADARS and *in vivo* expression tracking. a) Schematic of synthetic mRNA synthesis of RADARS with different base modifications. b) Comparison of different mRNA modifications for synthetic mRNA RADARS detecting IL6 transgene expression in HEK293FT cells when supplemented with MCP-ADAR2dd (E488Q, T490A) by plasmid transient transfection 24 hours before mRNA sensor transfection. c) Comparison of different mRNA modifications for synthetic mRNA RADARS detecting IL6 transgene expression in HEK293FT cells when supplemented with MCP-ADAR2dd (E488Q, T490A) mRNA at the time of sensor transfection. d) Comparison of mRNA SERPINA1 sensor activation fold change targeting different CCA sites on the SERPINA1 transcript in murine Hepa1-6 cells with transiently transfected tetracycline inducible human SERPINA1 expression. No exogenous ADAR is supplemented in this experiment. e) Comparison of Akaluc luminescence radiance in the NSG-PiZ mice and WT mice across non-targeting sensor, constitutive sensor, SERPINA1 CCA35 targeting sensor, and SERPINA1 CCA30 targeting sensor. Error bars indicate standard error of the mean as calculated by n=2 technical replicates.

**Extended Data Figure 10.**
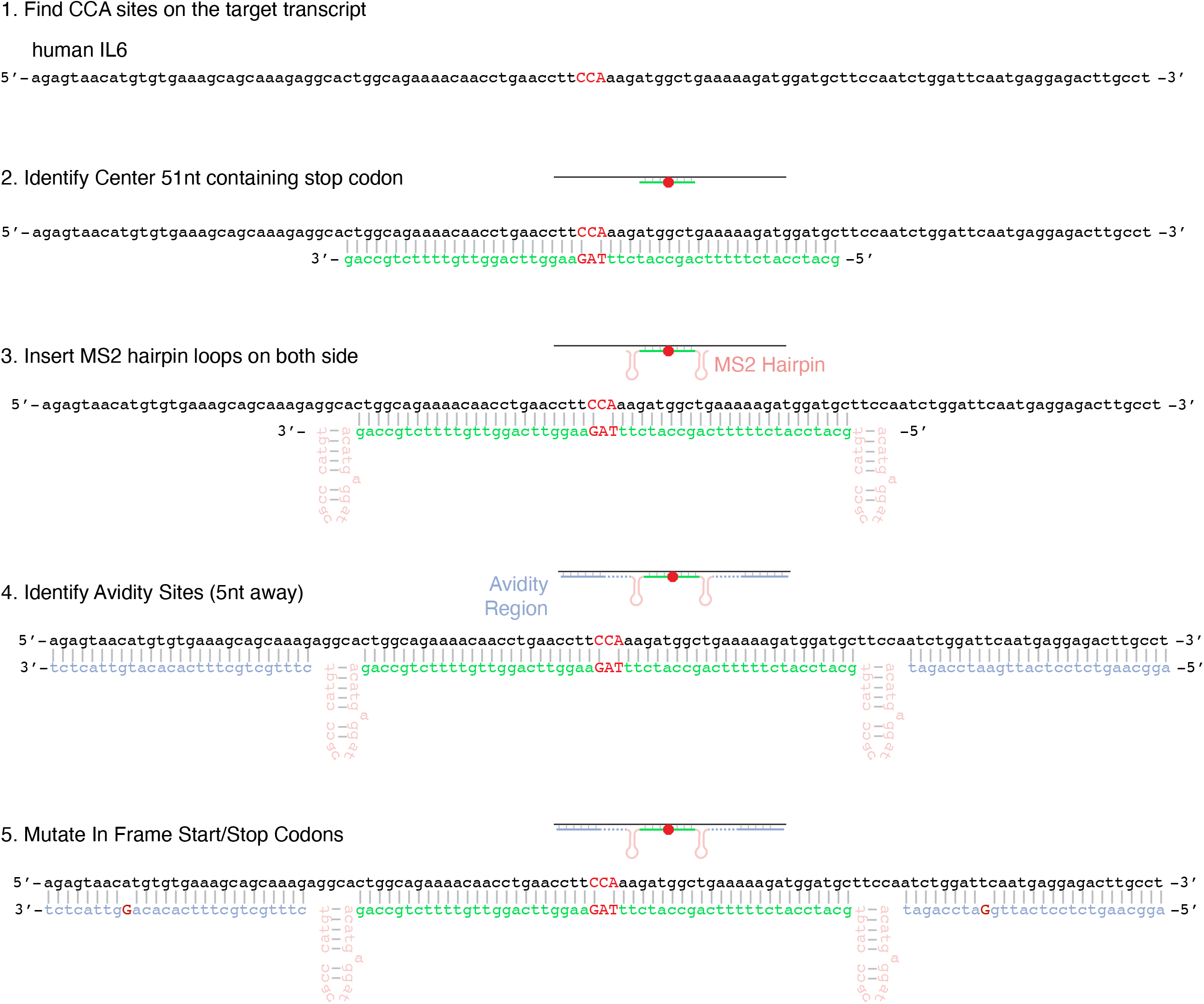
Schematic illustrating design of 3 site avidity binding RADARS.

## Methods

### Cloning of Luciferase Sensor

Luciferase sensors were cloned by Gibson assembly of PCR products. The sensor backbone is generated by cloning Cypridinia luciferase (Cluc) under expression of the CMV promoter and Gaussia luciferase (Gluc) under expression of the EF1-a promoter, both on a single vector. Expression of both luciferases on a single vector allowed one luciferase to serve as a dosing control for normalization of knockdown of the other luciferase, controlling for variation due to transfection conditions. Short sensors were ordered as primers and subsequently phosphorylated and annealed using T4 PNK. The annealed oligo is ligated into the backbone using T4 DNA Ligase (NEB) in a typical 10μL ligation reaction with 1μL of T4 DNA ligase, 30ng of the insert, 50ng of the backbone and 1μL of 10x ligation buffer at room temperature for 20 minutes. Long avidity sensor regions ordered as Eblocks directly from Integrated DNA Technologies (IDT). PCR products were purified by gel extraction (Monarch gel extraction kit, NEB), and assembled into the backbones using NEB HiFi DNA Assembly master mix kit, with 2.5μL of the mastermix, 30ng of backbone, and 5ng of the insert in a 5μL reaction. The reaction is incubated in the thermocycler at 50 degrees for 30 minutes and 2μL of assembled reactions were transformed into 20 μL of competent Stbl3 generated by Mix and Go! competency kit (Zymo) and plated on agar plates supplemented with appropriate antibiotics. After overnight growth at 37°C, colonies were picked into Terrific Broth (TB) media (Thermo Fisher Scientific) and incubated with shaking at 37°C for 24 hours. Cultures were harvested using QIAprep Spin Miniprep Kit (Qiagen) according to the manufacturer’s instructions.

### Cloning of Fluorescence RADARS

The fluorescence RADAR parent was cloned via Gibson assembly in three pieces, using pcDNA3.1(+) cut with HindIII and NotI as the backbone. mCherry was amplified off the Addgene vector 109427, and T2A mNeon was ordered as a gBlock from IDT. All fluorescence RADARS were subcloned into the parent fluorescence plasmid via golden gate cloning using the enzyme Esp3I (isoschizomer of BsmBI). Inserts were either ordered as complementary strands with overhangs and annealed with phosphorylation or produced via PCR. Golden-gate reactions used NEB BsmBIv2 golden gate assembly kit, or was assembled component-wise, in a 20 μL reaction containing 25 ng of vector and 2 μL of 1:200 diluted insert (approximately 5-10 ng). Reactions were thermocycled for 1 hour alternating between 25°C and 37°C for 5 minutes each, and then 0.75 μL of the reaction mix was transformed into 12.5 μL of Zymo Mix and Go competent cells. The transformed cells were diluted 1:1 with SOC media and 10 μL were streaked onto 50 ug/ mL carbenicillin agar plates. After incubation overnight at 37°C degrees, single colonies were picked into 4 mL of luria broth (LB) supplemented at 50 ug/ mL carbencillin. Plasmids were prepared from culture as described above for the luciferase sensors.

### Mammalian cell culture

HEK293FT cells (American Type Culture Collection (ATCC) - CRL32156), Hepa 1-6 cells (American Type Culture Collection (ATCC) -CRL-1830), and Hela cells (American Type Culture Collection (ATCC) - CCL-2) were cultured in Dulbecco’s Modified Eagle Medium with high glucose, sodium pyruvate, and GlutaMAX (Thermo Fisher Scientific), additionally supplemented with 10% (v/v) fetal bovine serum (FBS) and 1× penicillin-streptomycin (Thermo Fisher Scientific). For puromycin selection, HEK293FT cells were replated at a 1:3 dilution one day post-transfection into media supplemented with 1 μg/mL final concentration puromycin (Thermo Fisher Scientific). HEPG2 cells (American Type Culture Collection (ATCC – HB8065) were seeded in Eagle’s Minimum Essential Medium (Thermo Fisher Scientific), additionally supplemented with 10% (v/v) FBS, at 37°C and 5% CO2. Adherent cells were maintained at confluency below 80-90% at 37°C and 5% CO2.

### Transfection for luciferase sensors

Cells were plated at 5-15K the day prior to transfection in a 96-well plate coated with poly-D-lysine (BD Biocoat). HEK293FT and HepG2 cells were transfected with Lipofectamine 3000 (Thermo Fisher Scientific) according to manufacturer’s specifications. For all luciferase sensors, 20ng of ADAR plasmid, 20ng of sensor, and 20ng of targets were delivered to each well unless otherwise specified. For all mRNA transfection in the experiment, HEK293 cells were seeded at 15K the day prior to transfection, and 20ng of ADAR plasmid or pUC19 are co-transfected with 20ng of target transgene plasmid using Lipofectamine 3000. 24 hours later, the medium is removed and 50ng of the gaussia luciferase sensors mRNA mixed with the 50ng of cypridina luciferase mRNA in Lipofectamine MessengerMax (Thermo Fisher Scientific) following manufacturer’s specifications and transfected into each well.

### Measurement of luciferase activity

Media containing secreted luciferase was harvested 48 h after transfection, unless otherwise noted. 20μL of media is used to measure luciferase activity using Targeting Systems Cypridinia and Targeting systems Gaussia luciferase assay kits (Targeting Systems) on a Biotek Synergy 4 plate reader with an injection protocol. All replicates were performed as biological replicates.

### Transfection for fluorescence sensors

Cells were plated at 10K the day prior to transfection in Corning 96-well tissue-culture treated plates (black), resulting in approximately 40-50% confluency the day of transfection. For all fluorescence sensors, HEK293FT cells were transfected with 100 ng total plasmid DNA using TransIT-LT1 according to manufacturer specifications (ratio 1 ug DNA: 3 μL Trans reagent). Unless otherwise specified, RADARS, ADAR, and target plasmid were mixed at equal concentrations (33.3 ng / condition); for experiments without one or more of the previous, pUC19 was substituted accordingly to keep the total concentration of DNA at 100 ng.

### Confocal microscopy of fluorescent RADARS

48 hours post-transfection, all wells were measured via confocal microscopy under the following settings. For each well, a 2×2 image at 10x magnification was collected and stitched around the center point. Images were collected in 488 nm (32.8% power, 100 ms exposure), 561 nm (35.2% power, 100 ms exposure), 640 nm (80% power, 100 ms exposure), and brightfield channels (25 ms exposure).

### Quantification of fluorescence signal from images

Images were opened in Matlab, and segmented via watershed in the mCherry channel. For each segmented cell, the total pixel area and mean intensity of the pixels was computed for mNeon (488 nm), mCherry (561 nm), and iRFP (640 nm) channels, and output into an aggregated csv file. Csv files were batch processed in R with the following steps: all csv files were merged, conditions with low aggregated area (few cells, or not transfected with sensor) were merged, fluorescence background for each channel was subtracted from all conditions in that channel, and aggregated values for each condition were divided by area to obtain average fluorescence intensity. Standard deviation was computed by comparing average values in three technical transfection replicates. For mNeon/mCherry ratio values, the average mNeon fluorescence intensity for a condition was divided by the average mCherry value for that same condition. For fluorescence ratio values and ratio fold change values, error was propagated according to the formula:

### Quantification of percent mNeon positive cells in confocal images

We observed some consistent leakiness of the mNeon in the fluorescent sensors, either due to a very low level of plasmid contamination or ribosome slippage. Therefore, we gated the detection of mNeon positive cells at 30 AU above background, and determined the percent of mCherry positive cells in a condition that expressed mNeon higher than this threshold. mNeon values were plotted in log base 10 as a histogram with kernel density smoothing to generate the plots in Figure 2E.

### Extraction of RNA and next generation sequencing of RADARS

For calculating the editing rate of RADARS sensors, cells were harvested 48 hours post-transfection, after imaging. Total RNA was extracted using the RNeasy 96 Kit (Qiagen) with DNase treatment. cDNA was prepared with SuperScript IV reverse transcriptase (Invitrogen) and a sensor-specific primer. The guide regions of the sensors were amplified, indexed, and sequenced on an Illumina MiSeq platform. Reads were demultiplexed and aligned to each sensor, and the A-to-I editing rates were calculated with an in-house MATLAB pipeline.

### Quantification of protein expression

Two days after the transfection of HEK293FT cells, the Nano-Glo HiBiT Lytic Detection System (Promega) was used for the quantification of the HiBiT tags, in cell lysates. For the preparation of the Nano-Glo HiBiT Lytic Reagent, the Nano-Glo HiBit Lytic Buffer (Promega) was mixed with Nano-Glo HiBiT Lytic Substrate (Promega) and the LgBiT Protein (Promega) according to manufacturer’s protocol. The volume of Nano-Glo HiBiT Lytic Reagent added was equal to the culture medium present in each well, and the samples were placed on an orbital shaker at 600 rpm for 3 minutes. After incubation of 10 minutes at room temperature, the readout took place with 125 gain and 2 seconds integration time using a plate reader (Biotek Synergy Neo 2). The control background was subtracted from the final measurements.

### mRNA Synthesis

Before in vitro transcription, the DNA template was obtained by PCR with targeted forward primers containing T7 promoters. The sensor mRNA and the MCP-ADAR2dd mRNA were transcribed, and poly-A tailed using the HiScribe™ T7 ARCA mRNA Kit (NEB, E2065S) with 50% supplement of 5-Methyl-CTP and Pseudo-UTP (Jena Biosciences), following the manufacturer’s protocol. The mRNA was then cleaned up using the MEGAclear™ Transcription Clean-Up Kit (Thermo Fisher, AM1908).

### siRNA transfection

NEFM siRNA (Horizon Discovery, L-019968-01-0005), PPIB siRNA (Horizon Discovery, D-001820-01-05), or non-targeted control (NTC) siRNA (Thermo Fisher Scientific, AM4611) was transfected using the TransIT-TKO (Mirus Bio) at a final concentration of 100 nM. RADARS sensors were transfected 24 hours after siRNA transfection, and cells were assayed 24 hours after sensor transfection. The non-targeting sensor background was subtracted from the final measurements.

### Heat shock testing

HeLa cells (ATCC CCL-2) were transfected with either HSP40 or HSP70 RADARS. 24 hours post transfection, fresh media is supplemented to all cells and then a portion of cells are moved to 42 degree Celsius (5% CO2) for 24 hours. Media is harvested at the end of 24 hours of heat shock and subjected to luciferase measurements.

### Harvest of total RNA and quantitative PCR

For gene expression experiments in mammalian cells, cell harvesting and reverse transcription for cDNA generation was performed using a previously described modification of the commercial Cells-to-Ct kit (Thermo Fisher Scientific) 48 h after transfection(*1*). Transcript expression was then quantified with qPCR using Fast Advanced Master Mix (Thermo Fisher Scientific) and TaqMan qPCR probes (Thermo Fisher Scientific) with GAPDH control probes (Thermo Fisher Scientific). All qPCR reactions were performed in 10-μl reactions with two technical replicates in a 384-well format and read out using a LightCycler 480 Instrument II (Roche). For multiplexed targeting reactions, readout of different targets was performed in separate wells. Expression levels were calculated by subtracting housekeeping control (GAPDH) cycle threshold (Ct) values from target Ct values to normalize for total input, resulting in ΔCt levels. Relative transcript abundance was computed as 2−ΔCt. All replicates were performed as biological replicates.

### Cell Viability Assay

Mammalian cells were transfected with RADARS caspase, target, and MCP-ADAR2dd. Twenty-four hours after transfection, cells were split 1:5 into fresh media and the +drug samples were supplemented with 10nM of AP20187 (Sigma Aldrich). After 24 hours of additional growth, cells were assayed for viability by CellTiter-Glo Luminescent Cell Viability Assay (Promega).

### Computational design of avidity sensors

Avidity sensors are generated using python scripts in the following repository. (https://github.com/abugoot-lab/RADARS) Schematic for generation of a typical three avidity guide RADARS with two MS2 hairpin loops and 5nt spacing between the guide regions (on the target) are shown in Extended Data Figure 10.

### *In vivo* mRNA delivery and comparative in vivo bioluminescence

Prior to bioluminescence imaging, 8 to 10 week old Albino B6 and NSG-PiZ mice were anesthetized with 3% isoflurane and injected with 5 ug of synthesized mRNA via retro-orbital injection using in vivo-jetRNA transfection reagent (Polyplus). At 18 hours post-injection, the mice were anesthetized again with 3% isoflurane and immediately administered 100 μl of 15 mM AkaLumine-HCL (Sigma Aldrich) for imaging. Ventral bioluminescence images were acquired using an IVIS Spectrum In Vivo Imaging System (PerkinElmer). The following conditions were used for image acquisition: exposure time = 60 sec, binning = medium: 4, field of view = 12.5 × 12.5 cm, and f/stop = 1. Bioluminescent images were analyzed using Living Image 4.3 software (PerkinElmer).

### Animal husbandry and animal protocol

All experiments were carried out on female B6(Cg)-Tyr^c-2J^/J (Albino B6) and NOD.Cg-Prkdc^scid^ Il2rg^tm1Wjl^ Tg(SERPINA1*E342K)#Slcw/SzJ (NSG-PiZ) (The Jackson Laboratory) mice with ad libitum access to food and water. NSG-PiZ mice express the mutant human SERPINA1 on the immunodeficient NOD scid gamma background. All mice were housed in individually ventilated cages, in a temperature-controlled animal facility (normal 12:12 hour light-dark cycles) and used in accordance with approved procedures by the Committee on Animal Care at MIT.

## References

1. Rurik, J. G. et al. CAR T cells produced in vivo to treat cardiac injury. Science 375, 91–96 (2022).

2. Ronald, J. A., Chuang, H.-Y., Dragulescu-Andrasi, A., Hori, S. S. & Gambhir, S. S. Detecting cancers through tumor-activatable minicircles that lead to a detectable blood biomarker. Proc. Natl. Acad. Sci. U. S. A. 112, 3068–3073 (2015).

3. Rozenblatt-Rosen, O., Stubbington, M. J. T., Regev, A. & Teichmann, S. A. The Human Cell Atlas: from vision to reality. Nature 550, 451–453 (2017).

4. ENCODE Project Consortium. The ENCODE (ENCyclopedia Of DNA Elements) Project. Science 306, 636–640 (2004).

5. Mich, J. K. et al. Functional enhancer elements drive subclass-selective expression from mouse to primate neocortex. Cell Rep. 34, 108754 (2021).

6. Blankvoort, S., Witter, M. P., Noonan, J., Cotney, J. & Kentros, C. Marked Diversity of Unique Cortical Enhancers Enables Neuron-Specific Tools by Enhancer-Driven Gene Expression. Curr. Biol. 28, 2103–2114.e5 (2018).

7. Heinz, S., Romanoski, C. E., Benner, C. & Glass, C. K. The selection and function of cell type-specific enhancers. Nat. Rev. Mol. Cell Biol. 16, 144–154 (2015).

8. Nord, A. S. & West, A. E. Neurobiological functions of transcriptional enhancers. Nat. Neurosci. 23, 5–14 (2020).

9. Kim, J. et al. De novo-designed translation-repressing riboregulators for multi-input cellular logic. Nat. Chem. Biol. 15, 1173–1182 (2019).

10. Gambill, L., Staubus, A., Ameruoso, A. & Chappell, J. A split ribozyme that links detection of a native RNA to orthogonal protein outputs. bioRxiv 2022.01.12.476080 (2022) doi:10.1101/2022.01.12.476080.

11. Wang, S., Emery, N. J. & Liu, A. P. A Novel Synthetic Toehold Switch for MicroRNA Detection in Mammalian Cells. ACS Synth. Biol. 8, 1079–1088 (2019).

12. Li, Y., Teng, X., Zhang, K., Deng, R. & Li, J. RNA Strand Displacement Responsive CRISPR/Cas9 System for mRNA Sensing. Anal. Chem. 91, 3989–3996 (2019).

13. Hanewich-Hollatz, M. H., Chen, Z., Hochrein, L. M., Huang, J. & Pierce, N. A. Conditional Guide RNAs: Programmable Conditional Regulation of CRISPR/Cas Function in Bacterial and Mammalian Cells via Dynamic RNA Nanotechnology. ACS Cent Sci 5, 1241–1249 (2019).

14. Zhao, E. M. et al. RNA-responsive elements for eukaryotic translational control. Nat. Biotechnol. (2021) doi:10.1038/s41587-021-01068-2.

15. Stafforst, T. & Schneider, M. F. An RNA-Deaminase Conjugate Selectively Repairs Point Mutations. Angewandte Chemie International Edition vol. 51 11166–11169 (2012).

16. Montiel-Gonzalez, M. F., Vallecillo-Viejo, I., Yudowski, G. A. & Rosenthal, J. J. C. Correction of mutations within the cystic fibrosis transmembrane conductance regulator by site-directed RNA editing. Proceedings of the National Academy of Sciences 110, 18285–18290 (2013).

17. Cox, D. B. T. et al. RNA editing with CRISPR-Cas13. Science 358, 1019–1027 (2017).

18. Katrekar, D. et al. In vivo RNA editing of point mutations via RNA-guided adenosine deaminases. Nat. Methods 16, 239–242 (2019).

19. Qu, L. et al. Programmable RNA editing by recruiting endogenous ADAR using engineered RNAs. Nat. Biotechnol. 37, 1059–1069 (2019).

20. Reautschnig, P. et al. CLUSTER guide RNAs enable precise and efficient RNA editing with endogenous ADAR enzymes in vivo. Nat. Biotechnol. (2022) doi:10.1038/s41587-021-01105-0.

21. Abudayyeh, O. O. et al. A cytosine deaminase for programmable single-base RNA editing. Science 365, 382–386 (2019).

22. Bass, B. L. & Weintraub, H. An unwinding activity that covalently modifies its double-stranded RNA substrate. Cell 55, 1089–1098 (1988).

23. Wong, S. K., Sato, S. & Lazinski, D. W. Substrate recognition by ADAR1 and ADAR2. RNA 7, 846–858 (2001).

24. Kuttan, A. & Bass, B. L. Mechanistic insights into editing-site specificity of ADARs. Proc. Natl. Acad. Sci. U. S. A. 109, E3295–304 (2012).

25. Galipon, J., Ishii, R., Suzuki, Y., Tomita, M. & Ui-Tei, K. Differential Binding of Three Major Human ADAR Isoforms to Coding and Long Non-Coding Transcripts. Genes 8, (2017).

26. Merkle, T. et al. Precise RNA editing by recruiting endogenous ADARs with antisense oligonucleotides. Nat. Biotechnol. (2019) doi:10.1038/s41587-019-0013-6.

27. Matthews, M. M. et al. Structures of human ADAR2 bound to dsRNA reveal base-flipping mechanism and basis for site selectivity. Nat. Struct. Mol. Biol. 23, 426–433 (2016).

28. Uhlén, M. et al. Proteomics. Tissue-based map of the human proteome. Science 347, 1260419 (2015).

29. Chao, J. A., Patskovsky, Y., Almo, S. C. & Singer, R. H. Structural basis for the coevolution of a viral RNA-protein complex. Nat. Struct. Mol. Biol. 15, 103–105 (2008).

30. Straathof, K. C. et al. An inducible caspase 9 safety switch for T-cell therapy. Blood 105, 4247–4254 (2005).

31. Schwinn, M. K. et al. CRISPR-Mediated Tagging of Endogenous Proteins with a Luminescent Peptide. ACS Chem. Biol. 13, 467–474 (2018).

32. Litke, J. L. & Jaffrey, S. R. Highly efficient expression of circular RNA aptamers in cells using autocatalytic transcripts. Nat. Biotechnol. 37, 667–675 (2019).

33. Abe, N. et al. Rolling Circle Translation of Circular RNA in Living Human Cells. Sci. Rep. 5, 16435 (2015).

34. Boëlle, P.-Y., Debray, D., Guillot, L., Corvol, H. & French CF Modifier Gene Study Investigators. SERPINA1 Z allele is associated with cystic fibrosis liver disease. Genet. Med. 21, 2151–2155 (2019).

35. GTEx Consortium. The Genotype-Tissue Expression (GTEx) project. Nat. Genet. 45, 580–585 (2013).

36. Kauffman, K. J. et al. Efficacy and immunogenicity of unmodified and pseudouridine-modified mRNA delivered systemically with lipid nanoparticles in vivo. Biomaterials 109, 78–87 (2016).

37. Yeh, H.-W. et al. ATP-Independent Bioluminescent Reporter Variants To Improve in Vivo Imaging. ACS Chem. Biol. 14, 959–965 (2019).

