## Supplementary Information for "Programmable eukaryotic protein expression with RNA sensors"

[1] McGovern Institute for Brain Research at MIT
Massachusetts Institute of Technology
Cambridge, MA 02139, USA
[2] Department of Biological Engineering
Massachusetts Institute of Technology
Cambridge, MA 02139, USA
[3] Broad Institute of Harvard and MIT
Cambridge, MA 02142, USA
[4] Stem Cell and Regenerative Biology
Harvard University
Cambridge, MA 02139, USA
[5]Department of Systems, Synthetic, and Quantitative Biology
Harvard Medical School
Boston, MA 02115, USA

*These authors contributed equally.
‡These authors jointly supervised the work

### Supplementary Table

**Supplementary Table 1. Sequences of sensors used in this study**

| Description | Sequence (5'-3') |
|---|---|
| Non-Target | taccccgtacgacccgaatggtAgacgctaggctcagttttccacatctgg |
| IL6 51bp | gcatccatctttttcagccatctttAgaaggttcaggttgttttctgccag |
| IL6 81bp | gaatccagattggaagcatccatctttttcagccatctttAgaaggttcaggttgttttctgccagtgcctctttgctgct |
| IL6 171bp | ctccaaaagaccagtgatgattttcaccaggcaagtctcctcattgaatccagattggaagcatccatctttttcagccatctttAgaaggttcaggttgttttctgccagtgcctctttgctgctttcacacatgttactcttgttacatgtctcctttctcagggctgg |
| IL6 225bp | ctggaggtactctgggtatacctcaaactccaaaagaccagtgatgattttcaccaggcaagtctcctcattgaatccagattggaagcatccatctttttcagccatctttAgaaggttcaggttgttttctgccagtgcctctttgctgctttcacacatgttactcttgttacatgtctcctttctcagggctgggatgccgtcgaggatgtaccgaatttg |
| IL6 279bp | ttgttcctcactactctcaaatctgttctggaggtactctgggtatacctcaaactccaaaagaccagtgatgattttcaccaggcaagtctcctcattgaatccagattggaagcatccatctttttcagccatctttAgaaggttcaggttgttttctgccagtgcctctttgctgctttcacacatgttactcttgttacatgtctcctttctcagggctgggatgccgtcgaggatgtaccgaatttgtttgtcaattcgttctggagaggtgag |
| IL6 Three Avidity | atgggcaagtctcctcattggatccagatacatgaggatcacccatgtgcatccatctttttcagccatctttAgaaggttcaggttgttttctgccagacatggggatcacccaggtctttgctgctttcacacaggttactc |
| IL6 Five Avidity Far Spacing | ttgcacagctctggcttgttcctcactactctcaaatctgttctggaggtactacatgaggatcacccatgtggcaagtctcctcattggatccagatacatgaggatcacccatgtgcatccatctttttcagccatctttAgaaggttcaggttgttttctgccagacatggggatcacccaggtctttgctgctttcacacaggttactcacatggggatcacccaggtgggggtactggggcagggaaggcagcaggcaacaccaggagcagccccag |
| IL6 Five Avidity Middle Spacing | aagagccctcaggctggactgcaggaactccttaaagctgcgcagaatgagatacatgaggatcacccatgtcagctctggcttgttcctcactactctcaaatctgttctggaggtactctaggacatgaggatcacccatgtgcatccatctttttcagccatctttAgaaggttcaggttgttttctgccagacatggggatcacccaggtgctgggatgccgtcgaggatgtaccgaatttgtttgtcaattcgttctggacatggggatcacccaggtgggcagccccagggagaaggcaactggaccgaaggcgcttgtggagaagg |
| IL6 Five Avidity Close Spacing | actccaaaagaccagtggtggttttcgacgcaggaccaccgcgtcggcaagtctcctcattggatccagatacatgaggatcacccatgtgcatccatctttttcagccatctttAgaaggttcaggttgttttctgccaggacgcaggaccaccgcgtcctttgctgctttcacacaggttactcacatggggatcacccaggtacaggtctcctttctcagggctgaga |
| IL6 Seven Avidity | atctgttctggaggtactctaggtataaagggtggaggaacaccccaccctaactccaaaagaccagtgaggattttcacgggtggaggatcaccccacccgaggcaagtctcctcattgaatccagatacacatgaggatcacccatgtgcatccatctttttcagccatctttggaaggttcaggttgttttctgccagacagaagcaccatcagggcttctgctttgctgctttcacacatgttactctgtgcgtggagcatcagcccacgcacatgtctcctttctcagggctgagaggtcgcgaagagcatcagccttcgcgtcgttctggagaggtgagtAgctgtct |
| IL6 Seven Avidity Dual Stop Codon | atctgttctggaggtactctaggtataaagggtggaggaacaccccaccctaactccaaaagaccagtgaggattttcacgggtggaggatcaccccacccgaggcaagtctcctcattgaatccagatacacatgaggatcacccatgtgcatccatctttttcagccatctttAgaaggttcaggttgttttctgccagacagaagcaccatcagggcttctgctttgctgctttcacacatgttactctgtgcgtggagcatcagcccacgcacatgtctcctttctcagggctgagaggtcgcgaagagcatcagccttcgcgtcgttctggagaggtgagtAgctgtct |

| | |
|---|---|
| EGFP 51bp | tcagggtggtcacgagggtaggccagggcacgggcagcttgccggtggtgc |
| EGFP Five Avidity | tggtcgcgcttctcgttggggtctttgagacatgaggatcacccatgtggcggactgggtgctcaggtagtggttaagggtggaggaacaccccaccctgcagcagcacggggccgtcgccgatAggggtgttctgctggtagtggtcggacagaagcaccatcagggcttctgtgcacgctgccgtcctcgaggttgtgggtgcgtggagcatcagcccacgcacttgaagttcaccttgatgccgttctt |
| EGFP_IL6_AND_Gate | tggtcgcgcttctcgttggggtctttgtcgcgaagagcatcagccttcgcgggcggactgggtgctcaggtagtggttaagggtggaggaacaccccaccctgcagcagcacggggccgtcgccgatAggggtgttctgctggtagtggtcggacagaagcaccatcagggcttctgtgcacgctgccgtcctcgaggttgtgggtgcgtggagcatcagcccacgcacttgaagttcaccttgatgccgttcttactccaaaagaccagtggtggttttcgacgcaggaccaccgcgtcggcaagtctcctcattggatccagatacatgaggatcacccatgtgcatccatctttttcagccatctttAgaaggttcaggttgttttctgccaggacgcaggaccaccgcgtcctttgctgctttcacacaggttactcacatggggatcacccaggtacaggtctcctttctcagggctgaga |
| HSP40 | caggaacttttcgccgcatgccaggcctcgacgcaggaccaccgcgtcacatctttgaatacgacgggtatcgtcagacatgaggatcacccatgtgtccagagtggggacgttcactgtgcaaagggtggaggaacaccccaccctacagagcctcccggaggctggtcctAgcaggataaatggcatcagagccatacagaagcaccatcagggcttctgttaaagatattgtggggcttgtcctttgtgcgtggagcatcagcccacgcaaaagacgatatcagctggaatgttgttagcgcagaggaacaccctgcgtctggtctccttccttggggaaagtgg |
| HSP70 | tgcacctcgtcctccgctttgtacttcagacatgaggatcacccatgtctcctgcaccatgcgctcgatctcctcaagggtggaggaacaccccacccttcaggcggcccttgtcgttggtggtAgtggtcttgttggccttgccggtgcacagaagcaccatcagggcttctgtccgtggccgtgacgttcaggaggccggtgcgtggagcatcagcccacgcaatcgatgtcgaaggtcacctcgatctg |
| SERPINA1_CCA_2_five_avidity | gtatgggctgaaggcgaactcagccagacgggtggaggatcaccccacccggggtggtcttgttggaggttgggtgataagggtggaggaacaccccacccttcatggtgggatgtatctgtcttctAggcagcatctccctggggatcctcaacagaagcaccatcagggcttctgggagacagggaccaggcagcacaggccgtgcgtggagcatcagcccacgcagcaggaggatgccccacgagacagaag |
| SERPINA1_CCA_4_five_avidity | ggactggtgtgccagctggcggtatggacgggtggaggatcaccccacccgaggcgaactcagccaggttgggggtggaagggtggaggaacaccccaccctttgaaggttgggtggtcctggtcatAgtgggatgtatctgtcttctgggcaacagaagcaccatcagggcttctgtccctggggatcctcagccagggagacgtgcgtggagcatcagcccacgcaccaggcagcacaggcctgccagcagga |
| SERPINA1_CCA_5_five_avidity | agatattggtgctgttggactggtgtgacgggtggaggatcaccccacccgtggcggtataggctgaaggcgaactcaaagggtggaggaacaccccaccctgttgggggtgatcttgttgaaggttAggtgatcctgatcagggtgggatgtacagaagcaccatcagggcttctgtcttctgggcagcatctccctggggatgtgcgtggagcatcagcccacgcagccagggagacagggaccaggcagcac |
| SERPINA1_CCA_6_five_avidity | gcgatgctcactggggagaagaagataacgggtggaggatcaccccacccggctgttggactggtgtgccagctggcgaagggtggaggaacaccccaccctggctggaggcgaactcagccaggttAggggtggtcttgttggaggttgggtacagaagcaccatcagggcttctgtggtcatggtgggatgtatctgtcttcgtgcgtggagcatcagcccacgcaagcatctccctggggatcctcagccag |
| SERPINA1_CCA_7_fiv | cttggtccccagggagagcattgcaaaacgggtggaggatcaccccacccgtggcgatgctcactggggagaagaagaaagggtggaggaacaccccaccctgtgctgttggactggtgtgccagctAgcggtataggctgaaggcgaactcaacagaagcaccatcagggcttctggttgggggtgatcttgttgaaggttgggtgcgtggagcatcagcccacgcacctgatcatggtgggaggtatctgtct |

| | |
|---|---|
| e_avidity | |
| SERPINA1_CCA_8_five_avidity | gtgagtgtcagccttggtccccagggaacgggtggaggatcaccccacccgttgcaaaggctgtggcgatgctcactgaagggtggaggaacaccccaccctaagaagatattggtgctgttggactAgtgtgccagctggcggtataggctgacagaagcaccatcagggcttctggaactcagccaggttgggggtgatcttgtgcgtggagcatcagcccacgcaaggttgggtgatcctgatcatggtggg |
| SERPINA1_CCA_17_five_avidity | gtacaactttttaacatcctccaaaaaacgggtggaggatcaccccacccgccactagcttcaggccctcgctggggaaagggtggaggaacaccccaccctccattgccggtggtcagctggagctAgctgtctggctggttgagggtacggacagaagcaccatcagggcttctgttcctggaagccttcagggatctgagcgtgcgtggagcatcagcccacgcagaatctccgtggggttggaattcaggc |
| SERPINA1_CCA_18_five_avidity | tgggtggtacaactttttaacatcctcacgggtggaggatcaccccacccgacttatccactagcttcaggccctcgcaagggtggaggaacaccccaccctaacaggccattgccggtggtcagctAgagctggctgtctggctggttgaggacagaagcaccatcagggcttctggaggagttcctggaagccttcagggatgtgcgtggagcatcagcccacgcacctccggaatctccgtggggttggaat |
| SERPINA1_CCA_19_five_avidity | aaggcttctgagtggtacaactttttaacgggtggaggatcaccccacccgctccaaaaacttatccactggcttcagaagggtggaggaacaccccaccctcgctggggaacaggccattgccggtAgtcagctggagctggctgtctggctacagaagcaccatcagggcttctgagggtacggaggagttcctggaagcctgtgcgtggagcatcagcccacgcagatctgagcctccggaatctccgtgag |
| SERPINA1_CCA_20_five_avidity | tttcccttgagtacccttctccacgtaacgggtggaggatcaccccacccgtggtctgtttcttggcctcttcggtgtaagggtggaggaacaccccaccctaagttgacagtgaaggcttctgagtAgtacaactttttaacatcctccaaaacagaagcaccatcagggcttctgatccactggcttcaggccctcgctgaggtgcgtggagcatcagcccacgcaggccattgccggtggtcagctggagct |
| SERPINA1_CCA_22_five_avidity | ccagctggacagcttcttacagtgctgacgggtggaggatcaccccacccgtacacatgcctaaacgcttcatcatggaagggtggaggaacaccccaccctttcacggtggtcacctggtccacgtAgaagtcctcttcctcggtgtccttgacagaagcaccatcagggcttctgaaagggtctctcccatttgcctttaaagtgcgtggagcatcagcccacgcatgtaattcaccagagcaaaaactgtgt |
| SERPINA1_CCA_23_five_avidity | cagcagcacccagctggacagcttcttacgggtggaggatcaccccacccggctggatgttacacatgcctaaacgctaagggtggaggaacaccccaccctataggcaccttcacggtggtcacctAgtccacgtggaagtcctcttcctcgacagaagcaccatcagggcttctgcttgacttcaaagggtctctcccatttgtgcgtggagcatcagcccacgcatacagaagatgtaattcaccagagcaa |
| SERPINA1_CCA_24_five_avidity | tatttcatcagcagcacccagctggacacgggtggaggatcaccccacccgcttacagtgctggatgttaaacatgccaagggtggaggaacaccccaccctgcttcatcatgggcaccttcacggtAgtcacctggtccacgtggaagtcctacagaagcaccatcagggcttctgtcggtgtccttgacttcaaagggtctcgtgcgtggagcatcagcccacgcatttgcctttaaagaagatgtaattcac |
| SERPINA1_CCA_25_fi | tttcccctcatcaggcaggaagaagatacgggtggaggatcaccccacccgtggcattgcccaggtatttcatcagcaaagggtggaggaacaccccaccctcagctggacagcttcttacagtgctAgaggttaaacaggcctaaacgcttcacagaagcaccatcagggcttctgaggcaccttcacggtggtcacctggtcgtgcgtggagcatcagcccacgcaggaagtcctcttcctcggtgtccttgg |

| | |
|---|---|
| ve_avidity | |
| SERPINA1_CCA_26_five_avidity | tcattttccaggtgctgtagtttccccacgggtggaggatcaccccacccgaggcaggaagaagatggcggtggcattaagggtggaggaacaccccaccctggtatttcatcagcagcacccagctAgacagcttcttacagtgctggatgtacagaagcaccatcagggcttctgaggcctaaacgcttcatcataggcaccgtgcgtggagcatcagcccacgcaggtggtcacctggtccacgtggaagtc |
| SERPINA1_CCA_27_five_avidity | ttttccaggaacttggtgatgatatcgacgggtggaggatcaccccacccggagttcattttccaggtgctgtggtttaagggtggaggaacaccccaccctcatcaggcaggaagaagatggcggtAgcattgcccaggtatttcatcagcaacagaagcaccatcagggcttctgcagctggacagcttcttacagtgctgggtgcgtggagcatcagcccacgcaaaacatgcctacacgcttcatcatagg |
| SERPINA1_CCA_28_five_avidity | tcttcattttccaggaacttggtgatgacgggtggaggatcaccccacccggtgggtgagttcattttccaggtgctgaagggtggaggaacaccccacccttcccctcatcaggcaggaagaagatAgcggtggcattgcccaggtatttcaacagaagcaccatcagggcttctgagcacccagctggacagcttcttacaggtgcgtggagcatcagcccacgcagatgttaaacatgcctacacgcttcat |
| SERPINA1_CCA_29_five_avidity | atcataggttccagtaatggacagtttacgggtggaggatcaccccacccgaatgtaagctggcagaccttctgtcttaagggtggaggaacaccccacccttccaggaacttggtgatgatatcgtAggtgagttcattttccaggtgctgtacagaagcaccatcagggcttctgcccctcatcaggcaggaagaagatggcgtgcgtggagcatcagcccacgcacattgcccaggtatttcatcagcagca |
| SERPINA1_CCA_31_five_avidity | ttgctgaagaccttagtgatgcccagtacgggtggaggatcaccccacccgcaggacgctcttcagatcataggttccaagggtggaggaacaccccacccttggacagtttgggtaaatgtaagctAgcagaccttctgtcttcattttccaacagaagcaccatcagggcttctgttggtgaggatatcgtgggtgagttcagtgcgtggagcatcagcccacgcacaggtgctgtggtttcccctcatcagg |
| SERPINA1_CCA_32_five_avidity | gagaggtcagccccattgctgaagaccacgggtggaggatcaccccacccggatgcccagttgacccaggacgctcttaagggtggaggaacaccccaccctcataggttccagtactggacagtttAggtaaatgtaagctggcagaccttcacagaagcaccatcagggcttctgtcattttccaggaacttggtgaggatagtgcgtggagcatcagcccacgcaggtgagttcattttccaggtgctgtgg |
| SERPINA1_CCA_34_five_avidity | atagacatgggtatggcctctaaaaacacgggtggaggatcaccccacccgcccagcagcttcagtccctttctcgtcaagggtggaggaacaccccacccttcagcacagccttatgcacggccttAgagagcttcaggggtgcctcctctgacagaagcaccatcagggcttctgccggagaggtcagccccattgctgaaggtgcgtggagcatcagcccacgcaagtgatgcccagttgacccaggacgct |
| SERPINA1_CCA_35_five_avidity | ggtttgttgaacttgacctcgggggggacgggtggaggatcaccccacccgcatgggtatggcctctacaaacatggcaagggtggaggaacaccccaccctcagcttcagtccctttctcgtcgatAgtcagcacagccttatgcacggcctacagaagcaccatcagggcttctgagcttcaggggtgcctcctctgtgaccgtgcgtggagcatcagcccacgcagaggtcagccccattgctgaagacctt |
| SERPINA1_CCA_37_fi | atgaagaggggagacttggtattttgtacgggtggaggatcaccccacccgcattacgaagacaaagggtttgttgaaaagggtggaggaacaccccaccctcctcgggggggatggacatgggtatAgcctctaaaaacatggccccagcagacagaagcaccatcagggcttctggtccctttctcgtcgagggtcagcacagtgcgtggagcatcagcccacgcaaggcacggccttggagagcttcagggg |

| | |
|---|---|
| ve_avidity | |
| SERPINA1_CCA_30_five_avidity | aggacgctcttcagatcatgggttccaacgggtggaggatcaccccacccgggacagtttgggtacatgtacgctggcaagggtggaggaacaccccaccctttctgtcttcattttccaggaacttAgtggtggtatcgtgggtgggttcatacagaagcaccatcagggcttctgaggtgctgtagtttcccctcatcaggcgtgcgtggagcatcagcccacgcagaagatggcggtggcattgcccaggta |
| SERPINA1_CCA_30_seven_avidity | tgctggagaccttggtggtgcccagttagacatgaggatcacccatgtaggacgctcttcagatcatgggttccaacgggtggaggatcaccccacccgggacagtttgggtacatgtacgctggcaagggtggaggaacaccccaccctttctgtcttcattttccaggaacttAgtggtggtatcgtgggtgggttcatacagaagcaccatcagggcttctgaggtgctgtagtttcccctcatcaggcgtgcgtggagcatcagcccacgcagaagatggcggtggcattgcccaggtatcgacgcaggaccaccgcgtctttcatcagcagcacccagctggacag |
| 69 bp iRFP | TGCGGGAGCCAGTGTTCCGCTCAGATCAATGGGTAGCCCAGCCCGTTCCAGCTCAATGATCAGTCCTCC |
| dCas9 | TCTTCAGTTTCTTGGACTTGCCCTTTTCCACTTTAGCCACCACCAGCACAGAATAGGCCACGGTGGGGC |
| 69 bp NPY | CTGCTGGCGCGTCCTCGCCCGGATTGTCCGGCTTaGAGGGGTACCCCTCAGCCAGAATGCCCAAACACA |
| 51 bp iRFP | CAGTGTTCCGCTCAGATCAATGGGTAGCCCAGCCCGTTCCAGCTCAATGAT |
| 111 bp iRFP | TGCAGTTCGAATTCGCTCCAGTGCGGGAGCCAGTGTTCCGCTCAGATCAATGGGTAGCCCAGCCCGTTCCAGCTCAATGATCAGTCCTCCTTCAGGTGGTCGATGCATCAG |
| 150 bp iRFP | GCACAGAGCCCTCAGGGATCCTGCAGTTCGAATTCGCTCCAGTGCGGGAGCCAGTGTTCCGCTCAGATCAATGGGTAGCCCAGCCCGTTCCAGCTCAATGATCAGTCCTCCTTCAGGTGGTCGATGCATCAGTCCATCGTACTCTGTGGA |
| 201 bp iRFP | CTGGAACAGCAGTGCAGTATCGTCGCACAGAGCCCTCAGGGATCCTGCAGTTCGAATTCGCTCCAGTGCGGGAGCCAGTGTTCCGCTCAGATCAATGGGTaGCCCAGCCCGTTCCAGCTCAATGATCAGTCCTCCTTCAGGTGGTCGATCCATCAGTCCATCGTACTCTGTGGAGGGGTTTCCAATGCGGCACCTGACGGC |
| 249 bp iRFP | CACTCGGTCGTAGCCGGTACACTGCTGGAACAGCAGTGCAGTATCGTCGCACAGAGCCCTCAGGGATCCTGCAGTTCGAATTCGCTCCAGTGCGGGAGCCAGTGTTCCGCTCAGATCAATGGGTAGCCCAGCCCGTTCCAGCTCAATGATCAGTCCTCCTTCAGGTGGTCGATCCATCAGTCCATCGTACTCTGTGGAGGGGTTTCCAATGCGGCACCTGACGGCCACTGGCATTCCTTCTGCGGTAGG |
| 351 bp iRFP | TATTTCTGCATACACCTCTCCGTGCCCCTGTTCATCGAACCGATAGACCATCACTCGGTCGTAGCCGGTACACTGCTGGAACAGCAGTGCAGTATCGTCGCACAGAGCCCTCAGGGATCCTGCAGTTCGAATTCGCTCCAGTGCGGGAGCCAGTGTTCCGCTCAGATCAATGGGTAGCCCAGCCCGTTCCAGCTCAATGATCAGTCCTCCTTCAGGTGGTCGATCCATCAGTCCATCGTACTCTGTGGAGGGGTTTCCAATGCGGCACCTGACGGCCACTGGCATTCCTTCTGCGGTAGGGTCCAGGTGGGGCAGAATCTTGATCAGCAGATCCCCGTCGATCTCAGCCAG |

| | |
|---|---|
| 450 bp iRFP | GGAGCTAGGATACCTGTTGCCAAAGTAGCTCTCCAGTCCAGTCACATGT ATTTCTGCATACACCTCTCCGTGCCCCTGTTCATCGAACCGATAGACCA TCACTCGGTCGTAGCCGGTACACTGCTGGAACAGCAGTGCAGTATCGT CGCACAGAGCCCTCAGGGATCCTGCAGTTCGAATTCGCTCCAGTGCGG GAGCCAGTGTTCCGCTCAGATCAATGGGTAGCCCAGCCCGTTCCAGCTC AATGATCAGTCCTCCTTCAGGTGGTCGATCCATCAGTCCATCGTACTCT GTGGAGGGGTTTCCAATGCGGCACCTGACGGCCACTGGCATTCCTTCTG CGGTAGGGTCCAGGTGGGGCAGAATCTTGATCAGCAGATCCCCGTCGA TCTCAGCCAGGGGGACTCCCAGCACACTTCCCAGATTCAGGAATTCAG CGGCGTTGGCAGA |
| 600 bp iRFP | AGGCTGATAAGAGACATCCACCAGGACGCGCACCCTCTGTCTTTCGTA CAGTCTCCGGGCCATCTGTGGCACGAGGGAGCTAGGATACCTGTTGCC AAAGTAGCTCTCCAGTCCAGTCACATGTATTTCTGCATACACCTCTCCG TGCCCCTGTTCATCGAACCGATAGACCATCACTCGGTCGTAGCCGGTAC ACTGCTGGAACAGCAGTGCAGTATCGTCGCACAGAGCCCTCAGGGATC CTGCAGTTCGAATTCGCTCCAGTGCGGGAGCCAGTGTTCCGCTCAGATC AATGGGTAGCCCAGCCCGTTCCAGCTCAATGATCAGTCCTCCTTCAGGT GGTCGATCCATCAGTCCATCGTACTCTGTGGAGGGGTTTCCAATGCGGC ACCTGACGGCCACTGGCATTCCTTCTGCGGTAGGGTCCAGGTGGGGCA GAATCTTGATCAGCAGATCCCCGTCGATCTCAGCCAGGGGGACTCCCA GCACACTTCCCAGATTCAGGAATTCAGCGGCGTTGGCAGATGCCTGAA TGATTCTATCGTCAGGCTCGGAGACCACCAGCAGTGTTCCGTGTGGCTG GATGGATCCGGCCAGATC |
| 69 bp RPS5 | CGGCATACCGCCCTGCACTGTGAGGCAGGTACTTAGCATACTTCTCCTT CACTGCAATGTAATCCTGCA |
| 69 bp UBC | CACCCCCCCTCAAGCGCAGGACCAAGTGCAGAGTAGACTCTTTCTGGA TGTTGTAGTCAGACAGGGTGC |
| NEFM | TTTCTTTTCCTCTTTAGCTTCGGCTTCCTCTCCTTCAGCTTCAGCTTCTGT TTCTCCTTCTTCCTGCTCACCTTCCTCTTTTTCACTAGAGCCTTCCTTCTC GGATCCTCCCTCTTCGGCTTGGTCTGTCTTAGCTCCCTCATCTTCTTCCT CCTCTTCTTCCTGGCCTTCTTCTTCCTCCTTTTCCCCTTCCTCTTCTTTAA CTTCAGGTGCAGTTGCTTTCACTAGAGACTTTTTGGCAGCTACTTCTTCT TCTTCAGCTTCGGGTTCCTCTTCCTTTTCTTCTGCTGCTTCTTTCTTCTCT TCCTTCATGGAAACGGCCAATTCCTCTGTAATGGCTGTCAGGGCCTCTT CCATTTCTGTCTTCTCATCCTCCACTTTGGTTTCCTCTATGATCTCCTCG ACAAATTTGTGTTGGACCTTAAGCTTGGGAGCTTCCACCTTGGGTTT |
| PPIB | TTTGCCTGCGTTGGCCATGCTCACCCAGCCAGGCCCGTAGTGCTTCAGT TTGAAGTTCTCATCGGGGAAGCGCTCACCGTAGATGCTCTTTCCTCCTG TGCCATCTCCCCTGGTGAAGTCTCCGCCCTGGATCATGAAGTCCTTGAT TACACGTTGGAATTTGCTGTTTTTGTAGCCAAATCCTTTCTCTCCTGTAG CTATGGCCACAAAATTATCCACTGTTTTTAGAACAGTCTTTCCGAAGAG ACCAAAGATCACCCGGCCTACATCTTCATCTCCAATTCGTTGGTCAAAA TACACCTTGACGGTGACTTTGGGCCCCTTCTTCTTCTCATCGGCCGCAG AAGGTCCCGGCAGCAGCAGGAAGAAGACGGACCCCGCGATGAGGGCG |

| | |
|---|---|
| | GCGGCAAGGAGCACCTTCATGTTGCGTTCGGAGAGGCGCAGCATCCAC AGGCGGAGGCG |

**Supplementary Table 2.Protein sequences of ADAR variants**

| | |
|---|---|
| ADAR2 Full Length (E488Q) | MDIEDEENMSSSSTDVKENRNLDNVSPKDGSTPGPGEGSQLSNGGGGGPGRKRPLEEGSNGHSKYRLKKRRKTPGPVLPKNALMQLNEIKPGLQYTLLSQTGPVHAPLFVMSVEVNGQVFEGSGPTKKKAKLHAAEKALRSFVQFPNASEAHLAMGRTLSVNTDFTSDQADFPDTLFNGFETPDKAEPPFYVGSNGDDSFSSSGDLSLSASPVPASLAQPPLPVLPPFPPPSGKNPVMILNELRPGLKYDFLSESGESHAKSFVMSVVVDGQFFEGSGRNKKLAKARAAQSALAAIFNLHLDQTPSRQPIPSEGLQLHLPQVLADAVSRLVLGKFGDLTDNFSSPHARRKVLAGVVMTTGTDVKDAKVISVSTGTKCINGEYMSDRGLALNDCHAEIISRRSLLRFLYTQLELYLNNKDDQKRSIFQKSERGGFRLKENVQFHLYISTSPCGDARIFSPHEPILEEPADRHPNRKARGQLRTKIESGQGTIPVRSNASIQTWDGVLQGERLLTMSCSDKIARWNVVGIQGSLLSIFVEPIYFSSIILGSLYHGDHLSRAMYQRISNIEDLPPLYTLNKPLLSGISNAEARQPGKAPNFSVNWTVGDSAIEVINATTGKDELGRASRLCKHALYCRWMRVHGKVPSHLLRSKITKPNVYHESKLAAKEYQAAKARLFTAFIKAGLGAWVEKPTEQDQFSLTP* |
| ADAR2 V1 (#145-701) | MFPNASEAHLAMGRTLSVNTDFTSDQADFPDTLFNGFETPDKAEPPFYVGSNGDDSFSSSGDLSLSASPVPASLAQPPLPVLPPFPPPSGKNPVMILNELRPGLKYDFLSESGESHAKSFVMSVVVDGQFFEGSGRNKKLAKARAAQSALAAIFNLHLDQTPSRQPIPSEGLQLHLPQVLADAVSRLVLGKFGDLTDNFSSPHARRKVLAGVVMTTGTDVKDAKVISVSTGTKCINGEYMSDRGLALNDCHAEIISRRSLLRFLYTQLELYLNNKDDQKRSIFQKSERGGFRLKENVQFHLYISTSPCGDARIFSPHEPILEEPADRHPNRKARGQLRTKIESGQGTIPVRSNASIQTWDGVLQGERLLTMSCSDKIARWNVVGIQGSLLSIFVEPIYFSSIILGSLYHGDHLSRAMYQRISNIEDLPPLYTLNKPLLSGISNAEARQPGKAPNFSVNWTVGDSAIEVINATTGKDELGRASRLCKHALYCRWMRVHGKVPSHLLRSKITKPNVYHESKLAAKEYQAAKARLFTAFIKAGLGAWVEKPTEQDQFSLTP* |
| ADAR2 V2 (#231-701) | MPSGKNPVMILNELRPGLKYDFLSESGESHAKSFVMSVVVDGQFFEGSGRNKKLAKARAAQSALAAIFNLHLDQTPSRQPIPSEGLQLHLPQVLADAVSRLVLGKFGDLTDNFSSPHARRKVLAGVVMTTGTDVKDAKVISVSTGTKCINGEYMSDRGLALNDCHAEIISRRSLLRFLYTQLELYLNNKDDQKRSIFQKSERGGFRLKENVQFHLYISTSPCGDARIFSPHEPILEEPADRHPNRKARGQLRTKIESGQGTIPVRSNASIQTWDGVLQGERLLTMSCSDKIARWNVVGIQGSLLSIFVEPIYFSSIILGSLYHGDHLSRAMYQRISNIEDLPPLYTLNKPLLSGISNAEARQPGKAPNFSVNWTVGDSAIEVINATTGKDELGRASRLCKHALYCRWMRVHGKVPSHLLRSKITKPNVYHESKLAAKEYQAAKARLFTAFIKAGLGAWVEKPTEQDQFSLTP* |
| ADAR2 V3 (#299-701) | MLHLDQTPSRQPIPSEGLQLHLPQVLADAVSRLVLGKFGDLTDNFSSPHARRKVLAGVVMTTGTDVKDAKVISVSTGTKCINGEYMSDRGLALNDCHAEIISRRSLLRFLYTQLELYLNNKDDQKRSIFQKSERGGFRLKENVQFHLYISTSPCGDARIFSPHEPILEEPADRHPNRKARGQLRTKIESGQGTIPVRSNASIQTWDGVLQGERLLTMSCSDKIARWNVVGIQGSLLSIFVEPIYFSSIILGSLYHGDHLSRAMYQRISNIEDLPPLYTLNKPLLSGISNAEARQPGKAPNFSVNWTVGDSAIEVINATTGKDELGRASRLCKHALYCRWMRVHGKVPSHLLRSKITKPNVYHESKLAAKEYQAAKARLFTAFIKAGLGAWVEKPTEQDQFSLTP* |
| ADAR | MVKDAKVISVSTGTKCINGEYMSDRGLALNDCHAEIISRRSLLRFLYTQLELYLNNKDDQKRSIFQKSERGGFRLKENVQFHLYISTSPCGDARIFSPHEPILEEPADR |

| | |
|---|---|
| 2 V4 (#363-701) | HPNRKARGQLRTKIESGQGTIPVRSNASIQTWDGVLQGERLLTMSCSDKIARWNVVGIQGSLLSIFVEPIYFSSIILGSLYHGDHLSRAMYQRISNIEDLPPLYTLNKPLLSGISNAEARQPGKAPNFSVNWTVGDSAIEVINATTGKDELGRASRLCKHALYCRWMRVHGKVPSHLLRSKITKPNVYHESKLAAKEYQAAKARLFTAFIKAGLGAWVEKPTEQDQFSLTP* |
| ADAR2 V5 (#395-701) | MAEIISRRSLLRFLYTQLELYLNNKDDQKRSIFQKSERGGFRLKENVQFHLYISTSPCGDARIFSPHEPILEEPADRHPNRKARGQLRTKIESGQGTIPVRSNASIQTWDGVLQGERLLTMSCSDKIARWNVVGIQGSLLSIFVEPIYFSSIILGSLYHGDHLSRAMYQRISNIEDLPPLYTLNKPLLSGISNAEARQPGKAPNFSVNWTVGDSAIEVINATTGKDELGRASRLCKHALYCRWMRVHGKVPSHLLRSKITKPNVYHESKLAAKEYQAAKARLFTAFIKAGLGAWVEKPTEQDQFSLTP* |
| ADAR2 V6 (#452-701) | MGDARIFSPHEPILEEPADRHPNRKARGQLRTKIESGQGTIPVRSNASIQTWDGVLQGERLLTMSCSDKIARWNVVGIQGSLLSIFVEPIYFSSIILGSLYHGDHLSRAMYQRISNIEDLPPLYTLNKPLLSGISNAEARQPGKAPNFSVNWTVGDSAIEVINATTGKDELGRASRLCKHALYCRWMRVHGKVPSHLLRSKITKPNVYHESKLAAKEYQAAKARLFTAFIKAGLGAWVEKPTEQDQFSLTP* |
| MCP-ADAR2dd (E488Q/T490A) | MGPKKKRKVGVKMASNFTQFVLVDNGGTGDVTVAPSNFANGIAEWISSNSRSQAYKVTCSVRQSSAQNRKYTIKVEVPKGAWRSYLNMELTIPIFATNSDCELIVKAMQGLLKDGNPIPSAIAANSGIYAMASNFTQFVLVDNGGTGDVTVAPSNFANGIAEWISSNSRSQAYKVTCSVRQSSAQNRKYTIKVEVPKGAWRSYLNMELTIPIFATNSDCELIVKAMQGLLKDGNPIPSAIAANSGGGSGGTGGSGGTQLHLPQVLADAVSRLVLGKFGDLTDNFSSPHARRKVLAGVVMTTGTDVKDAKVISVSTGTKCINGEYMSDRGLALNDCHAEIISRRSLLRFLYTQLELYLNNKDDQKRSIFQKSERGGFRLKENVQFHLYISTSPCGDARIFSPHEPILEEPADRHPNRKARGQLRTKIESGQGAIPVRSNASIQTWDGVLQGERLLTMSCSDKIARWNVVGIQGSLLSIFVEPIYFSSIILGSLYHGDHLSRAMYQRISNIEDLPPLYTLNKPLLSGISNAEARQPGKAPNFSVNWTVGDSAIEVINATTGKDELGRASRLCKHALYCRWMRVHGKVPSHLLRSKITKPNVYHESKLAAKEYQAAKARLFTAFIKAGLGAWVEKPTEQDQFSLT* |
| flADAR1 p150 | MKLMNPRQGYSLSGYYTHPFQGYEHRQLRYQQPGPGSSPSSFLLKQIEFLKGQLPEAPVIGKQTPSLPPSLPGLRPRFPVLLASSTRGRQVDIRGVPRGVHLGSQGLQRGFQHPSPRGRSLPQRGVDCLSSHFQELSIYQDQEQRILKFLEELGEGKATTAHDLSGKLGTPKKEINRVLYSLAKKGKLQKEAGTPPLWKIAVSTQAWNQHSGVVRPDGHSQGAPNSDPSLEPEDRNSTSVSEDLLEPFIAVSAQAWNQHSGVVRPDSHSQGSPNSDPGLEPEDSNSTSALEDPLEFLDMAEIKEKICDYLFNVSDSSALNLAKNIGLTKARDINAVLIDMERQGDVYRQGTTPPIWHLTDKKRERMQIKRNTNSVPETAPAAIPETKRNAEFLTCNIPTSNASNNMVTTEKVENGQEPVIKLENRQEARPEPARLKPPVHYNGPSKAGYVDFENGQWATDDIPDDLNSIRAAPGEFRAIMEMPSFYSHGLPRCSPYKKLTECQLKNPISGLLEYAQFASQTCEFNMIEQSGPPHEPRFKFQVVINGREFPPAEAGSKKVAKQDAAMKAMTILLEEAKAKDSGKSEESSHYSTEKESEKTAESQTPTPSATSFFSGKSPVTTLLECMHKLGNSCEFRLLSKEGPAHEP |

| | |
|---|---|
| | KFQYCVAVGAQTFPSVSAPSKKVAKQMAAEEAMKALHGEATNSMASDNQPE<br>GMISESLDNLESMMPNKVRKIGELVRYLNTNPVGGLLEYARSHGFAAEFKLVD<br>QSGPPHEPKFVYQAKVGGRWFPAVCAHSKKQGKQEAADAALRVLIGENEKAE<br>RMGFTEVTPVTGASLRRTMLLLSRSPEAQPKTLPLTGSTFHDQIAMLSHRCFNT<br>LTNSFQPSLLGRKILAAIIMKKDSEDMGVVVSLGTGNRCVKGDSLSLKGETVN<br>DCHAEIISRRGFIRFLYSELMKYNSQTAKDSIFEPAKGGEKLQIKKTVSFHLYIST<br>APCGDGALFDKSCSDRAMESTESRHYPVFENPKQGKLRTKVENGEGTIPVESSD<br>IVPTWDGIRLGERLRTMSCSDKILRWNVLGLQGALLTHFLQPIYLKSVTLGYLF<br>SQGHLTRAICCRVTRDGSAFEDGLRHPFIVNHPKVGRVSIYDSKRQSGKTKETS<br>VNWCLADGYDLEILDGTRGTVDGPRNELSRVSKKNIFLLFKKLCSFRYRRDLL<br>RLSYGEAKKAARDYETAKNYFKKGLKDMGYGNWISKPQEEKNFYLCPV |
| flADAR1p110 | MKLMAEIKEKICDYLFNVSDSSALNLAKNIGLTKARDINAVLIDMERQGDVYR<br>QGTTPPIWHLTDKKRERMQIKRNTNSVPETAPAAIPETKRNAEFLTCNIPTSNAS<br>NNMVTTEKVENGQEPVIKLENRQEARPEPARLKPPVHYNGPSKAGYVDFENG<br>QWATDDIPDDLNSIRAAPGEFRAIMEMPSFYSHGLPRCSPYKKLTECQLKNPISG<br>LLEYAQFASQTCEFNMIEQSGPPHEPRFKFQVVINGREFPPAEAGSKKVAKQDA<br>AMKAMTILLEEAKAKDSGKSEESSHYSTEKESEKTAESQTPTPSATSFFSGKSPV<br>TTLLECMHKLGNSCEFRLLSKEGPAHEPKFQYCVAVGAQTFPSVSAPSKKVAK<br>QMAAEEAMKALHGEATNSMASDNQPEGMISESLDNLESMMPNKVRKIGELVR<br>YLNTNPVGGLLEYARSHGFAAEFKLVDQSGPPHEPKFVYQAKVGGRWFPAVC<br>AHSKKQGKQEAADAALRVLIGENEKAERMGFTEVTPVTGASLRRTMLLLSRSP<br>EAQPKTLPLTGSTFHDQIAMLSHRCFNTLTNSFQPSLLGRKILAAIIMKKDSEDM<br>GVVVSLGTGNRCVKGDSLSLKGETVNDCHAEIISRRGFIRFLYSELMKYNSQTA<br>KDSIFEPAKGGEKLQIKKTVSFHLYISTAPCGDGALFDKSCSDRAMESTESRHYP<br>VFENPKQGKLRTKVENGEGTIPVESSDIVPTWDGIRLGERLRTMSCSDKILRWN<br>VLGLQGALLTHFLQPIYLKSVTLGYLFSQGHLTRAICCRVTRDGSAFEDGLRHP<br>FIVNHPKVGRVSIYDSKRQSGKTKETSVNWCLADGYDLEILDGTRGTVDGPRN<br>ELSRVSKKNIFLLFKKLCSFRYRRDLLRLSYGEAKKAARDYETAKNYFKKGLK<br>DMGYGNWISKPQEEKNFYLCPV |
| flADAR2R455G | MDIEDEENMSSSSTDVKENRNLDNVSPKDGSTPGPGEGSQLSNGGGGGPGRKR<br>PLEEGSNGHSKYRLKKRRKTPGPVLPKNALMQLNEIKPGLQYTLLSQTGPVHA<br>PLFVMSVEVNGQVFEGSGPTKKKAKLHAAEKALRSFVQFPNASEAHLAMGRT<br>LSVNTDFTSDQADFPDTLFNGFETPDKAEPPFYVGSNGDDSFSSSGDLSLSASPV<br>PASLAQPPLPVLPPFPPPSGKNPVMILNELRPGLKYDFLSESGESHAKSFVMSVV<br>VDGQFFEGSGRNKKLAKARAAQSALAAIFNLHLDQTPSRQPIPSEGLQLHLPQV<br>LADAVSRLVLGKFGDLTDNFSSPHARRKVLAGVVMTTGTDVKDAKVISVSTG<br>TKCINGEYMSDRGLALNDCHAEIISRRSLLRFLYTQLELYLNNKDDQKRSIFQK<br>SERGGFRLKENVQFHLYISTSPCGDAGIFSPHEPILEEPADRHPNRKARGQLRTKI<br>ESGEGTIPVRSNASIQTWDGVLQGERLLTMSCSDKIARWNVVGIQGSLLSIFVEP<br>IYFSSIILGSLYHGDHLSRAMYQRISNIEDLPPLYTLNKPLLSGISNAEARQPGKA<br>PNFSVNWTVGDSAIEVINATTGKDELGRASRLCKHALYCRWMRVHGKVPSHL<br>LRSKITKPNVYHESKLAAKEYQAAKARLFTAFIKAGLGAWVEKPTEQDQFSLT<br>P |
| flADAR2 | MDIEDEENMSSSSTDVKENRNLDNVSPKDGSTPGPGEGSQLSNGGGGGPGRKR<br>PLEEGSNGHSKYRLKKRRKTPGPVLPKNALMQLNEIKPGLQYTLLSQTGPVHA<br>PLFVMSVEVNGQVFEGSGPTKKKAKLHAAEKALRSFVQFPNASEAHLAMGRT |

| | |
|---|---|
| S48<br>6T | LSVNTDFTSDQADFPDTLFNGFETPDKAEPPFYVGSNGDDSFSSSGDLSLSASPV<br>PASLAQPPLPVLPPFPPPSGKNPVMILNELRPGLKYDFLSESGESHAKSFVMSVV<br>VDGQFFEGSGRNKKLAKARAAQSALAAIFNLHLDQTPSRQPIPSEGLQLHLPQV<br>LADAVSRLVLGKFGDLTDNFSSPHARRKVLAGVVMTTGTDVKDAKVISVSTG<br>TKCINGEYMSDRGLALNDCHAEIISRRSLLRFLYTQLELYLNNKDDQKRSIFQK<br>SERGGFRLKENVQFHLYISTSPCGDARIFSPHEPILEEPADRHPNRKARGQLRTKI<br>ETGEGTIPVRSNASIQTWDGVLQGERLLTMSCSDKIARWNVVGIQGSLLSIFVEP<br>IYFSSIILGSLYHGDHLSRAMYQRISNIEDLPPLYTLNKPLLSGISNAEARQPGKA<br>PNFSVNWTVGDSAIEVINATTGKDELGRASRLCKHALYCRWMRVHGKVPSHL<br>LRSKITKPNVYHESKLAAKEYQAAKARLFTAFIKAGLGAWVEKPTEQDQFSLT<br>P |
| flA<br>DA<br>R2<br>T37<br>5G<br>E48<br>8Q<br>T49<br>0A | MDIEDEENMSSSSTDVKENRNLDNVSPKDGSTPGPGEGSQLSNGGGGGPGRKR<br>PLEEGSNGHSKYRLKKRRKTPGPVLPKNALMQLNEIKPGLQYTLLSQTGPVHA<br>PLFVMSVEVNGQVFEGSGPTKKKAKLHAAEKALRSFVQFPNASEAHLAMGRT<br>LSVNTDFTSDQADFPDTLFNGFETPDKAEPPFYVGSNGDDSFSSSGDLSLSASPV<br>PASLAQPPLPVLPPFPPPSGKNPVMILNELRPGLKYDFLSESGESHAKSFVMSVV<br>VDGQFFEGSGRNKKLAKARAAQSALAAIFNLHLDQTPSRQPIPSEGLQLHLPQV<br>LADAVSRLVLGKFGDLTDNFSSPHARRKVLAGVVMTTGTDVKDAKVISVSTG<br>GKCINGEYMSDRGLALNDCHAEIISRRSLLRFLYTQLELYLNNKDDQKRSIFQK<br>SERGGFRLKENVQFHLYISTSPCGDARIFSPHEPILEEPADRHPNRKARGQLRTKI<br>ESGQGAIPVRSNASIQTWDGVLQGERLLTMSCSDKIARWNVVGIQGSLLSIFVE<br>PIYFSSIILGSLYHGDHLSRAMYQRISNIEDLPPLYTLNKPLLSGISNAEARQPGK<br>APNFSVNWTVGDSAIEVINATTGKDELGRASRLCKHALYCRWMRVHGKVPSH<br>LLRSKITKPNVYHESKLAAKEYQAAKARLFTAFIKAGLGAWVEKPTEQDQFSL<br>TP |
| flA<br>DA<br>R2<br>R47<br>4E | MDIEDEENMSSSSTDVKENRNLDNVSPKDGSTPGPGEGSQLSNGGGGGPGRKR<br>PLEEGSNGHSKYRLKKRRKTPGPVLPKNALMQLNEIKPGLQYTLLSQTGPVHA<br>PLFVMSVEVNGQVFEGSGPTKKKAKLHAAEKALRSFVQFPNASEAHLAMGRT<br>LSVNTDFTSDQADFPDTLFNGFETPDKAEPPFYVGSNGDDSFSSSGDLSLSASPV<br>PASLAQPPLPVLPPFPPPSGKNPVMILNELRPGLKYDFLSESGESHAKSFVMSVV<br>VDGQFFEGSGRNKKLAKARAAQSALAAIFNLHLDQTPSRQPIPSEGLQLHLPQV<br>LADAVSRLVLGKFGDLTDNFSSPHARRKVLAGVVMTTGTDVKDAKVISVSTG<br>TKCINGEYMSDRGLALNDCHAEIISRRSLLRFLYTQLELYLNNKDDQKRSIFQK<br>SERGGFRLKENVQFHLYISTSPCGDARIFSPHEPILEEPADRHPNEKARGQLRTKI<br>ESGEGTIPVRSNASIQTWDGVLQGERLLTMSCSDKIARWNVVGIQGSLLSIFVEP<br>IYFSSIILGSLYHGDHLSRAMYQRISNIEDLPPLYTLNKPLLSGISNAEARQPGKA<br>PNFSVNWTVGDSAIEVINATTGKDELGRASRLCKHALYCRWMRVHGKVPSHL<br>LRSKITKPNVYHESKLAAKEYQAAKARLFTAFIKAGLGAWVEKPTEQDQFSLT<br>P |
| flA<br>DA<br>R2<br>T37<br>5G<br>E48<br>8Q | MDIEDEENMSSSSTDVKENRNLDNVSPKDGSTPGPGEGSQLSNGGGGGPGRKR<br>PLEEGSNGHSKYRLKKRRKTPGPVLPKNALMQLNEIKPGLQYTLLSQTGPVHA<br>PLFVMSVEVNGQVFEGSGPTKKKAKLHAAEKALRSFVQFPNASEAHLAMGRT<br>LSVNTDFTSDQADFPDTLFNGFETPDKAEPPFYVGSNGDDSFSSSGDLSLSASPV<br>PASLAQPPLPVLPPFPPPSGKNPVMILNELRPGLKYDFLSESGESHAKSFVMSVV<br>VDGQFFEGSGRNKKLAKARAAQSALAAIFNLHLDQTPSRQPIPSEGLQLHLPQV<br>LADAVSRLVLGKFGDLTDNFSSPHARRKVLAGVVMTTGTDVKDAKVISVSTG |

| | |
|---|---|
| | GKCINGEYMSDRGLALNDCHAEIISRRSLLRFLYTQLELYLNNKDDQKRSIFQK SERGGFRLKENVQFHLYISTSPCGDARIFSPHEPILEEPADRHPNRKARGQLRTKI ESGQGTIPVRSNASIQTWDGVLQGERLLTMSCSDKIARWNVVGIQGSLLSIFVEP IYFSSIILGSLYHGDHLSRAMYQRISNIEDLPPLYTLNKPLLSGISNAEARQPGKA PNFSVNWTVGDSAIEVINATTGKDELGRASRLCKHALYCRWMRVHGKVPSHL LRSKITKPNVYHESKLAAKEYQAAKARLFTAFIKAGLGAWVEKPTEQDQFSLT P |
| flA DA R2 T37 5G | MDIEDEENMSSSSTDVKENRNLDNVSPKDGSTPGPGEGSQLSNGGGGGPGRKR PLEEGSNGHSKYRLKKRRKTPGPVLPKNALMQLNEIKPGLQYTLLSQTGPVHA PLFVMSVEVNGQVFEGSGPTKKKAKLHAAEKALRSFVQFPNASEAHLAMGRT LSVNTDFTSDQADFPDTLFNGFETPDKAEPPFYVGSNGDDSFSSSGDLSLSASPV PASLAQPPLPVLPPFPPPSGKNPVMILNELRPGLKYDFLSESGESHAKSFVMSVV VDGQFFEGSGRNKKLAKARAAQSALAAIFNLHLDQTPSRQPIPSEGLQLHLPQV LADAVSRLVLGKFGDLTDNFSSPHARRKVLAGVVMTTGTDVKDAKVISVSTG GKCINGEYMSDRGLALNDCHAEIISRRSLLRFLYTQLELYLNNKDDQKRSIFQK SERGGFRLKENVQFHLYISTSPCGDARIFSPHEPILEEPADRHPNRKARGQLRTKI ESGEGTIPVRSNASIQTWDGVLQGERLLTMSCSDKIARWNVVGIQGSLLSIFVEP IYFSSIILGSLYHGDHLSRAMYQRISNIEDLPPLYTLNKPLLSGISNAEARQPGKA PNFSVNWTVGDSAIEVINATTGKDELGRASRLCKHALYCRWMRVHGKVPSHL LRSKITKPNVYHESKLAAKEYQAAKARLFTAFIKAGLGAWVEKPTEQDQFSLT P |
| flA DA R2 N47 3D | MDIEDEENMSSSSTDVKENRNLDNVSPKDGSTPGPGEGSQLSNGGGGGPGRKR PLEEGSNGHSKYRLKKRRKTPGPVLPKNALMQLNEIKPGLQYTLLSQTGPVHA PLFVMSVEVNGQVFEGSGPTKKKAKLHAAEKALRSFVQFPNASEAHLAMGRT LSVNTDFTSDQADFPDTLFNGFETPDKAEPPFYVGSNGDDSFSSSGDLSLSASPV PASLAQPPLPVLPPFPPPSGKNPVMILNELRPGLKYDFLSESGESHAKSFVMSVV VDGQFFEGSGRNKKLAKARAAQSALAAIFNLHLDQTPSRQPIPSEGLQLHLPQV LADAVSRLVLGKFGDLTDNFSSPHARRKVLAGVVMTTGTDVKDAKVISVSTG TKCINGEYMSDRGLALNDCHAEIISRRSLLRFLYTQLELYLNNKDDQKRSIFQK SERGGFRLKENVQFHLYISTSPCGDARIFSPHEPILEEPADRHPDRKARGQLRTKI ESGEGTIPVRSNASIQTWDGVLQGERLLTMSCSDKIARWNVVGIQGSLLSIFVEP IYFSSIILGSLYHGDHLSRAMYQRISNIEDLPPLYTLNKPLLSGISNAEARQPGKA PNFSVNWTVGDSAIEVINATTGKDELGRASRLCKHALYCRWMRVHGKVPSHL LRSKITKPNVYHESKLAAKEYQAAKARLFTAFIKAGLGAWVEKPTEQDQFSLT P |
| AD AR 2D D WT | MQLHLPQVLADAVSRLVLGKFGDLTDNFSSPHARRKVLAGVVMTTGTDVKD AKVISVSTGTKCINGEYMSDRGLALNDCHAEIISRRSLLRFLYTQLELYLNNKD DQKRSIFQKSERGGFRLKENVQFHLYISTSPCGDARIFSPHEPILEEPADRHPNRK ARGQLRTKIESGEGTIPVRSNASIQTWDGVLQGERLLTMSCSDKIARWNVVGIQ GSLLSIFVEPIYFSSIILGSLYHGDHLSRAMYQRISNIEDLPPLYTLNKPLLSGISNA EARQPGKAPNFSVNWTVGDSAIEVINATTGKDELGRASRLCKHALYCRWMRV HGKVPSHLLRSKITKPNVYHESKLAAKEYQAAKARLFTAFIKAGLGAWVEKPT EQDQFSLT |
| flA DA R2 | MDIEDEENMSSSSTDVKENRNLDNVSPKDGSTPGPGEGSQLSNGGGGGPGRKR PLEEGSNGHSKYRLKKRRKTPGPVLPKNALMQLNEIKPGLQYTLLSQTGPVHA PLFVMSVEVNGQVFEGSGPTKKKAKLHAAEKALRSFVQFPNASEAHLAMGRT |

| | |
|---|---|
| T490S | LSVNTDFTSDQADFPDTLFNGFETPDKAEPPFYVGSNGDDSFSSSGDLSLSASPV PASLAQPPLPVLPPFPPPSGKNPVMILNELRPGLKYDFLSESGESHAKSFVMSVV VDGQFFEGSGRNKKLAKARAAQSALAAIFNLHLDQTPSRQPIPSEGLQLHLPQV LADAVSRLVLGKFGDLTDNFSSPHARRKVLAGVVMTTGTDVKDAKVISVSTG TKCINGEYMSDRGLALNDCHAEIISRRSLLRFLYTQLELYLNNKDDQKRSIFQK SERGGFRLKENVQFHLYISTSPCGDARIFSPHEPILEEPADRHPNRKARGQLRTKI ESGEGSIPVRSNASIQTWDGVLQGERLLTMSCSDKIARWNVVGIQGSLLSIFVEP IYFSSIILGSLYHGDHLSRAMYQRISNIEDLPPLYTLNKPLLSGISNAEARQPGKA PNFSVNWTVGDSAIEVINATTGKDELGRASRLCKHALYCRWMRVHGKVPSHL LRSKITKPNVYHESKLAAKEYQAAKARLFTAFIKAGLGAWVEKPTEQDQFSLT P |
| flADAR2 WT | MDIEDEENMSSSSTDVKENRNLDNVSPKDGSTPGPGEGSQLSNGGGGGPGRKR PLEEGSNGHSKYRLKKRRKTPGPVLPKNALMQLNEIKPGLQYTLLSQTGPVHA PLFVMSVEVNGQVFEGSGPTKKKAKLHAAEKALRSFVQFPNASEAHLAMGRT LSVNTDFTSDQADFPDTLFNGFETPDKAEPPFYVGSNGDDSFSSSGDLSLSASPV PASLAQPPLPVLPPFPPPSGKNPVMILNELRPGLKYDFLSESGESHAKSFVMSVV VDGQFFEGSGRNKKLAKARAAQSALAAIFNLHLDQTPSRQPIPSEGLQLHLPQV LADAVSRLVLGKFGDLTDNFSSPHARRKVLAGVVMTTGTDVKDAKVISVSTG TKCINGEYMSDRGLALNDCHAEIISRRSLLRFLYTQLELYLNNKDDQKRSIFQK SERGGFRLKENVQFHLYISTSPCGDARIFSPHEPILEEPADRHPNRKARGQLRTKI ESGEGSIPVRSNASIQTWDGVLQGERLLTMSCSDKIARWNVVGIQGSLLSIFVEP IYFSSIILGSLYHGDHLSRAMYQRISNIEDLPPLYTLNKPLLSGISNAEARQPGKA PNFSVNWTVGDSAIEVINATTGKDELGRASRLCKHALYCRWMRVHGKVPSHL LRSKITKPNVYHESKLAAKEYQAAKARLFTAFIKAGLGAWVEKPTEQDQFSLT P |
| MCP-ADAR2DD WT | MGVKMASNFTQFVLVDNGGTGDVTVAPSNFANGIAEWISSNSRSQAYKVTCS VRQSSAQNRKYTIKVEVPKGAWRSYLNMELTIPIFATNSDCELIVKAMQGLLK DGNPIPSAIAANSGIYAMASNFTQFVLVDNGGTGDVTVAPSNFANGIAEWISSN SRSQAYKVTCSVRQSSAQNRKYTIKVEVPKGAWRSYLNMELTIPIFATNSDCEL IVKAMQGLLKDGNPIPSAIAANSGGGSGGTGGSGGTQLHLPQVLADAVSRLVL GKFGDLTDNFSSPHARRKVLAGVVMTTGTDVKDAKVISVSTGTKCINGEYMS DRGLALNDCHAEIISRRSLLRFLYTQLELYLNNKDDQKRSIFQKSERGGFRLKE NVQFHLYISTSPCGDARIFSPHEPILEEPADRHPNRKARGQLRTKIESGEGTIPVR SNASIQTWDGVLQGERLLTMSCSDKIARWNVVGIQGSLLSIFVEPIYFSSIILGSL YHGDHLSRAMYQRISNIEDLPPLYTLNKPLLSGISNAEARQPGKAPNFSVNWTV GDSAIEVINATTGKDELGRASRLCKHALYCRWMRVHGKVPSHLLRSKITKPNV YHESKLAAKEYQAAKARLFTAFIKAGLGAWVEKPTEQDQFSLT |
| flADAR2 R455E | MDIEDEENMSSSSTDVKENRNLDNVSPKDGSTPGPGEGSQLSNGGGGGPGRKR PLEEGSNGHSKYRLKKRRKTPGPVLPKNALMQLNEIKPGLQYTLLSQTGPVHA PLFVMSVEVNGQVFEGSGPTKKKAKLHAAEKALRSFVQFPNASEAHLAMGRT LSVNTDFTSDQADFPDTLFNGFETPDKAEPPFYVGSNGDDSFSSSGDLSLSASPV PASLAQPPLPVLPPFPPPSGKNPVMILNELRPGLKYDFLSESGESHAKSFVMSVV VDGQFFEGSGRNKKLAKARAAQSALAAIFNLHLDQTPSRQPIPSEGLQLHLPQV LADAVSRLVLGKFGDLTDNFSSPHARRKVLAGVVMTTGTDVKDAKVISVSTG TKCINGEYMSDRGLALNDCHAEIISRRSLLRFLYTQLELYLNNKDDQKRSIFQK SERGGFRLKENVQFHLYISTSPCGDAEIFSPHEPILEEPADRHPNRKARGQLRTKI |

| | |
|---|---|
| | ESGEGTIPVRSNASIQTWDGVLQGERLLTMSCSDKIARWNVVGIQGSLLSIFVEP IYFSSIILGSLYHGDHLSRAMYQRISNIEDLPPLYTLNKPLLSGISNAEARQPGKA PNFSVNWTVGDSAIEVINATTGKDELGRASRLCKHALYCRWMRVHGKVPSHL LRSKITKPNVYHESKLAAKEYQAAKARLFTAFIKAGLGAWVEKPTEQDQFSLT P |
| flA DA R2 T37 5G T49 0A | MDIEDEENMSSSSTDVKENRNLDNVSPKDGSTPGPGEGSQLSNGGGGGPGRKR PLEEGSNGHSKYRLKKRRKTPGPVLPKNALMQLNEIKPGLQYTLLSQTGPVHA PLFVMSVEVNGQVFEGSGPTKKKAKLHAAEKALRSFVQFPNASEAHLAMGRT LSVNTDFTSDQADFPDTLFNGFETPDKAEPPFYVGSNGDDSFSSSGDLSLSASPV PASLAQPPLPVLPPFPPPSGKNPVMILNELRPGLKYDFLSESGESHAKSFVMSVV VDGQFFEGSGRNKKLAKARAAQSALAAIFNLHLDQTPSRQPIPSEGLQLHLPQV LADAVSRLVLGKFGDLTDNFSSPHARRKVLAGVVMTTGTDVKDAKVISVSTG GKCINGEYMSDRGLALNDCHAEIISRRSLLRFLYTQLELYLNNKDDQKRSIFQK SERGGFRLKENVQFHLYISTSPCGDARIFSPHEPILEEPADRHPNRKARGQLRTKI ESGEGAIPVRSNASIQTWDGVLQGERLLTMSCSDKIARWNVVGIQGSLLSIFVEP IYFSSIILGSLYHGDHLSRAMYQRISNIEDLPPLYTLNKPLLSGISNAEARQPGKA PNFSVNWTVGDSAIEVINATTGKDELGRASRLCKHALYCRWMRVHGKVPSHL LRSKITKPNVYHESKLAAKEYQAAKARLFTAFIKAGLGAWVEKPTEQDQFSLT P |
| flA DA R2 R51 0E | MDIEDEENMSSSSTDVKENRNLDNVSPKDGSTPGPGEGSQLSNGGGGGPGRKR PLEEGSNGHSKYRLKKRRKTPGPVLPKNALMQLNEIKPGLQYTLLSQTGPVHA PLFVMSVEVNGQVFEGSGPTKKKAKLHAAEKALRSFVQFPNASEAHLAMGRT LSVNTDFTSDQADFPDTLFNGFETPDKAEPPFYVGSNGDDSFSSSGDLSLSASPV PASLAQPPLPVLPPFPPPSGKNPVMILNELRPGLKYDFLSESGESHAKSFVMSVV VDGQFFEGSGRNKKLAKARAAQSALAAIFNLHLDQTPSRQPIPSEGLQLHLPQV LADAVSRLVLGKFGDLTDNFSSPHARRKVLAGVVMTTGTDVKDAKVISVSTG TKCINGEYMSDRGLALNDCHAEIISRRSLLRFLYTQLELYLNNKDDQKRSIFQK SERGGFRLKENVQFHLYISTSPCGDARIFSPHEPILEEPADRHPNRKARGQLRTKI ESGEGTIPVRSNASIQTWDGVLQGEELLTMSCSDKIARWNVVGIQGSLLSIFVEP IYFSSIILGSLYHGDHLSRAMYQRISNIEDLPPLYTLNKPLLSGISNAEARQPGKA PNFSVNWTVGDSAIEVINATTGKDELGRASRLCKHALYCRWMRVHGKVPSHL LRSKITKPNVYHESKLAAKEYQAAKARLFTAFIKAGLGAWVEKPTEQDQFSLT P |
| flA DA R2 R45 5S | MDIEDEENMSSSSTDVKENRNLDNVSPKDGSTPGPGEGSQLSNGGGGGPGRKR PLEEGSNGHSKYRLKKRRKTPGPVLPKNALMQLNEIKPGLQYTLLSQTGPVHA PLFVMSVEVNGQVFEGSGPTKKKAKLHAAEKALRSFVQFPNASEAHLAMGRT LSVNTDFTSDQADFPDTLFNGFETPDKAEPPFYVGSNGDDSFSSSGDLSLSASPV PASLAQPPLPVLPPFPPPSGKNPVMILNELRPGLKYDFLSESGESHAKSFVMSVV VDGQFFEGSGRNKKLAKARAAQSALAAIFNLHLDQTPSRQPIPSEGLQLHLPQV LADAVSRLVLGKFGDLTDNFSSPHARRKVLAGVVMTTGTDVKDAKVISVSTG TKCINGEYMSDRGLALNDCHAEIISRRSLLRFLYTQLELYLNNKDDQKRSIFQK SERGGFRLKENVQFHLYISTSPCGDASIFSPHEPILEEPADRHPNRKARGQLRTKI ESGEGTIPVRSNASIQTWDGVLQGERLLTMSCSDKIARWNVVGIQGSLLSIFVEP IYFSSIILGSLYHGDHLSRAMYQRISNIEDLPPLYTLNKPLLSGISNAEARQPGKA PNFSVNWTVGDSAIEVINATTGKDELGRASRLCKHALYCRWMRVHGKVPSHL |

| | |
|---|---|
| | LRSKITKPNVYHESKLAAKEYQAAKARLFTAFIKAGLGAWVEKPTEQDQFSLT<br>P |
| flA<br>DA<br>R2<br>V35<br>1L | MDIEDEENMSSSSTDVKENRNLDNVSPKDGSTPGPGEGSQLSNGGGGGPGRKR<br>PLEEGSNGHSKYRLKKRRKTPGPVLPKNALMQLNEIKPGLQYTLLSQTGPVHA<br>PLFVMSVEVNGQVFEGSGPTKKKAKLHAAEKALRSFVQFPNASEAHLAMGRT<br>LSVNTDFTSDQADFPDTLFNGFETPDKAEPPFYVGSNGDDSFSSSGDLSLSASPV<br>PASLAQPPLPVLPPFPPPSGKNPVMILNELRPGLKYDFLSESGESHAKSFVMSVV<br>VDGQFFEGSGRNKKLAKARAAQSALAAIFNLHLDQTPSRQPIPSEGLQLHLPQV<br>LADAVSRLVLGKFGDLTDNFSSPHARRKLLAGVVMTTGTDVKDAKVISVSTGT<br>KCINGEYMSDRGLALNDCHAEIISRRSLLRFLYTQLELYLNNKDDQKRSIFQKS<br>ERGGFRLKENVQFHLYISTSPCGDARIFSPHEPILEEPADRHPNRKARGQLRTKIE<br>SGEGTIPVRSNASIQTWDGVLQGERLLTMSCSDKIARWNVVGIQGSLLSIFVEPI<br>YFSSIILGSLYHGDHLSRAMYQRISNIEDLPPLYTLNKPLLSGISNAEARQPGKAP<br>NFSVNWTVGDSAIEVINATTGKDELGRASRLCKHALYCRWMRVHGKVPSHLL<br>RSKITKPNVYHESKLAAKEYQAAKARLFTAFIKAGLGAWVEKPTEQDQFSLTP |

**Supplementary Table 3. NGS primers used for RNA editing analysis**

| | |
|---|---|
| EGFP_NGS_F | ACACTCTTTCCCTACACGACGCTCTTCCGATCTCcacggggccgtcgcc |
| EGFP_NGS_R | GTGACTGGAGTTCAGACGTGTGCTCTTCCGATCTcctgtgtgaaagcagcaaaggacgc |
| IL6_AviditySensors_NGS_F | ACACTCTTTCCCTACACGACGCTCTTCCGATCTCacatgaggatcacccatgtgcatcca |
| IL6_AviditySensors_NGS_R | GTGACTGGAGTTCAGACGTGTGCTCTTCCGATCTAGCAGGCTGAAGTTAGTAGCTCCg |
| SERPINA1_CCA30_NGS_F | ACACTCTTTCCCTACACGACGCTCTTCCGATCTCaacaccccaccctttctgtcttcattttcc |
| SERPINA1_CCA30_NGS_R | GTGACTGGAGTTCAGACGTGTGCTCTTCCGATCTAGCAGGCTGAAGTTAGTAGCTCCg |
| IL6_51bp_NGS_F | ACACTCTTTCCCTACACGACGCTCTTCCGATCTCCGGgatcctctagagccaccatg |
| IL6_51bp_NGS_R | GTGACTGGAGTTCAGACGTGTGCTCTTCCGATCTAGCAGGCTGAAGTTAGTAGCTCCg |
| FluorescentSensor_NGS_F | TCGTCGGCAGCGTCAGATGTGTATAAGAGACAGaacgaggactacaccatcgtg |
| FluorescentSensor_NGS_R | GTCTCGTGGGCTCGGAGATGTGTATAAGAGACAGCCCAGcccagtaccctgtccaacca |

**Supplementary Table 4: qPCR primers used in this study**

| | |
|---|---|
| IL6_qPCR_F | GGAGACTTGCCTGGTGAAA |
| IL6_qPCR_R | CTGGCTTGTTCCTCACTACTC |
| ACTB_qPCR_F | CATGTACGTTGCTATCCAGGC |
| ACTB_qPCR_R | CTCCTTAATGTCACGCACGAT |
| hsp40_qPCR_F | GAGTTCTTCGGTGGCAGAAA |
| hsp40_qPCR_R | CCATAGGGAAGCCAGAGAATG |
| hsp70_qPCR_F | GAGTCCTACGCCTTCAACAT |
| hsp70_qPCR_R | CTTGTCCAGCACCTTCTTCT |
| GAPDH_qPCR_F | ATTCCACCCATGGCAAATTC |
| GAPDH_qPCR_R | TGGGATTTCCATTGATGACAAG |
| NEFM_F | ACAACCACGACCTCAGCAGCTA |
| NEFM_R | GTTGAGGAGGTCCTGGTATTCG |
| PPIB_F | AACGCAGGCAAAGACACCAACG |
| PPIB_R | TCTGTCTTGGTGCTCTCCACCT |
| MDA5 (IFIH1)_F | aggaggaactgttgacaattg |
| MDA5 (IFIH1)_R | agtagctctcttacacctgattc |
| IFNB1_F | ttcagtgtcagaagctcctgtgg |
| IFNB1_R | ctgcttaatctcctcagggatgtca |
